# Biomechanical filtering supports tactile encoding efficiency in the human hand

**DOI:** 10.1101/2023.11.10.565040

**Authors:** Neeli Tummala, Gregory Reardon, Bharat Dandu, Yitian Shao, Hannes P. Saal, Yon Visell

## Abstract

Touching an object elicits skin oscillations that are biomechanically transmitted throughout the hand, driving responses in thousands of tactile receptors, including numerous exquisitely sensitive Pacinian corpuscles (PCs). Accepted descriptions of PC functionality characterize their response properties as highly stereotyped, based on experimental data gathered when stimuli are applied near the receptor. However, during natural touch, spiking activity in the majority of PCs is evoked by transmitted skin oscillations that are modified by biomechanical filtering. This filtering mechanism, stemming from dispersive wave dynamics in the skin, bears some similarity to the pre-neuronal filtering of auditory signals by the basilar membrane, a mechanical process that is instrumental to perception. Thus, we sought to clarify how skin biomechanics might influence tactile information encoding in the periphery. We used vibrometry imaging and computational neural experiments to examine the influence of biomechanical filtering on neural activity in whole-hand PC populations. We observed complex, location- and frequency-dependent patterns of filtering that were shaped by tissue mechanics and hand morphology. This source of biomechanical modulation diversified PC population spiking activity and enhanced tactile information encoding efficiency. These findings indicate that biomechanics furnishes a pre-neuronal mechanism that facilitates efficient tactile encoding and processing.

## INTRODUCTION

The sense of touch is stimulated when the skin comes into contact with the environment. During such contact events, perceptual information is often regarded as originating from the responses of tactile sensory neurons terminating near the contact location. But the sense of touch is also invoked when the environment is explored indirectly through a probe, such as a tool, fingernail, or whisker. Such probes are not innervated by sensory neurons. Instead, perceptual information is mediated by “internal contacts” that biomechanically couple the probe to skin innervated by tactile sensory neurons^1^. The same biomechanical couplings that mediate indirect touch are also involved during direct touch. In both cases, these couplings facilitate the transmission of touch-elicited skin oscillations to regions far from the contact location^2,3^, which excites remote tactile sensory neurons^4^.

Indeed, manual touch interactions, such as texture exploration^5,6^, dexterous manipulation^7^, and tool use^8^, generate prominent skin oscillations that are transmitted across the hand. The biomechanics of the hand transforms localized contact forces into spatially distributed skin oscillations that carry information about contact at the skin surface^6,9,10^. These oscillations excite widespread Pacinian corpuscle neurons (PCs)^3,11–13^ that encode the transmitted tactile information via spiking responses conveyed to the brain.

Central somatosensory processing integrates tactile inputs from receptors across the hand, reflecting a topographically complex somatotopic organization^14,15^. The central integration of widespread peripheral neural activity is, unsurprisingly, reflected in touch perception. For example, humans can discriminate between different surface textures^16^ or vibration frequencies^17^ applied to an anesthetized finger, utilizing information encoded by remote PCs. The perceptual significance of PC responses evoked across the hand is also exemplified by tactile summation, masking, and transfer learning effects observed when stimuli are applied at separate hand locations^18–20^. Across species, PCs also respond readily to subtle vibrations transmitted through the ground or other substrates, resulting from distant contact events^21–24^. This sensitivity enhances the ability of several species to detect vibrations that facilitate perception and communication over substantial distances^23,24^. In addition, PCs facilitate the perception of contact events during object manipulation and tool use, highlighting their integral role in both direct and mediated tactile perception^7,8,25^.

Recent studies suggest that hand biomechanics modifies transmitted skin oscillations through frequency-dependent and location-specific filtering imparted by tissues^11,26^. However, the specific effects of biomechanical filtering on PC spiking activity throughout the hand are unknown. Existing peripheral neural recordings reveal PC response characteristics to be highly stereotyped, with the highest frequency sensitivity between 200 to 300 Hz^27–31^. However, these recordings are only obtained from PCs closest to the stimulus contact location. As a result, they do not capture the effects of biomechanical filtering, which predominantly influence the responses of distant PCs. Thus, the influence of biomechanical filtering on tactile encoding by whole-hand PC populations and its implications for tactile sensing has received little prior attention.

It is not straightforward to deduce biomechanical influences on PC population responses due to the complex and heterogeneous morphology of the hand. Furthermore, extant experimental techniques preclude the simultaneous recording of neural signals from populations of PCs^32^. To overcome these limitations, we captured high-resolution vibrometry measurements of skin oscillations across the whole hand. We then used location-specific vibrometry measurements to selectively drive each receptor in a whole-hand population of spiking PC neuron models. We analyzed the vibrometry measurements to determine the frequency- and location-dependent patterns of biomechanical filtering. Using our data-driven neural simulation methodology, we then characterized the location-specific influences of biomechanical filtering on the tuning characteristics of PCs located throughout the hand, and the implications for tactile encoding by whole-hand PC populations. Our findings reveal that biomechanical filtering in the periphery furnishes a pre-neuronal mechanism that modulates and diversifies PC population spiking responses, thereby supporting efficient somatosensory encoding and processing.

## RESULTS

### Imaging whole-hand biomechanical transmission

We characterized the transmission of skin oscillations across the glabrous skin of several human hands (n = 7, P1 to P7). Mechanical impulses (0.5 ms duration) were applied at four distinct contact locations, and evoked skin oscillations were recorded at 200 to 350 spatially distributed locations via optical vibrometry (sample rate 20 kHz, grid spacing 8 mm; see Methods; Figure 1A). These impulse measurements characterized biomechanical transmission across the hand within the frequency range relevant to PCs (20 to 800 Hz). The dispersive nature of biomechanical transmission altered both the temporal structure and frequency content of skin oscillations (Figures 1B, 1D). As a consequence, we observed the pairwise temporal and spectral correlation of skin oscillations at different locations to decrease with increasing pairwise distance (Figures 1C, 1D).

**Figure 1.**
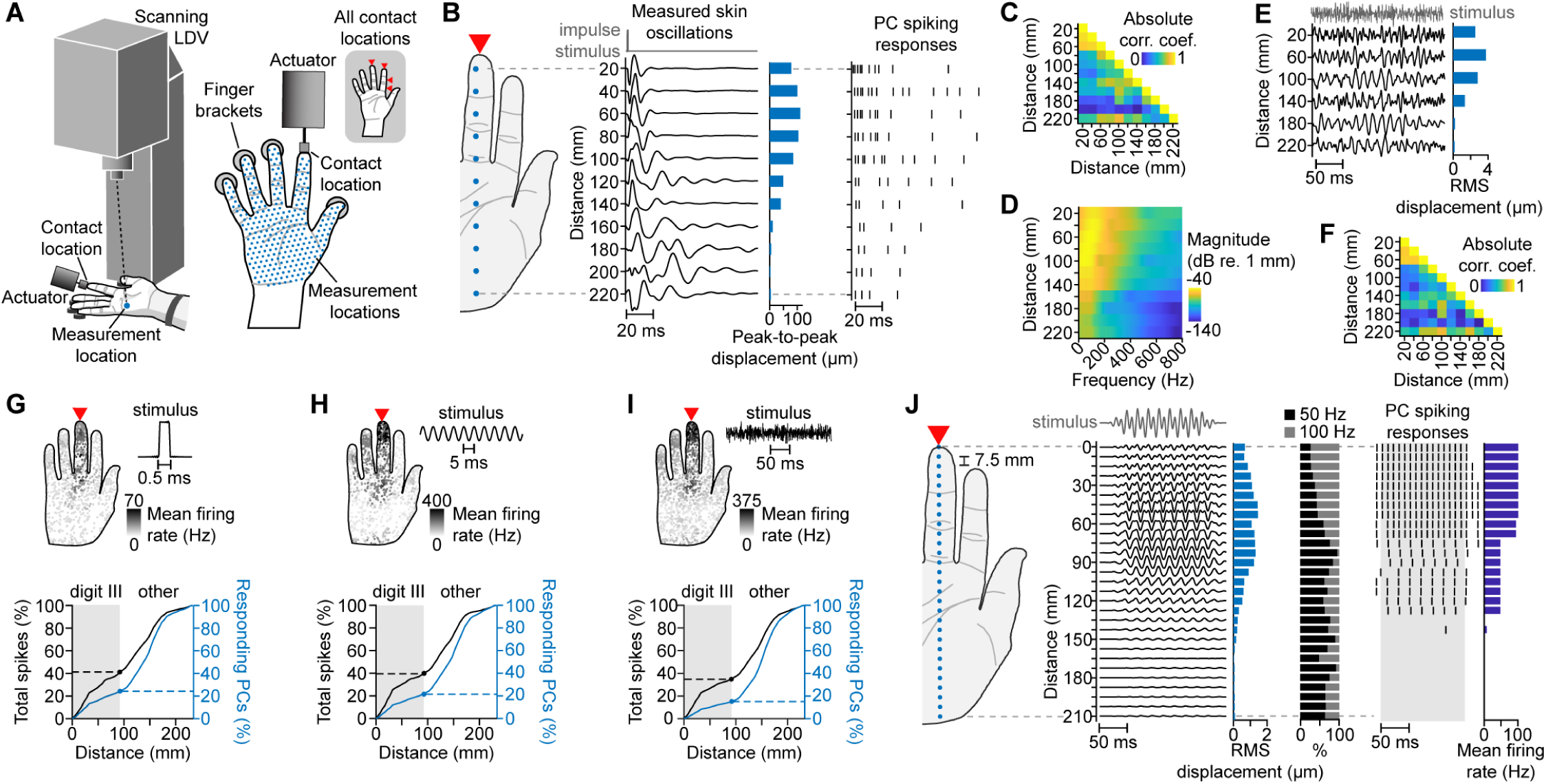
Evoked skin oscillations drive location-specific spiking responses in PCs throughout the hand. (A) Scanning laser Doppler vibrometer (LDV) measurement setup. (B) Left: vibrometry measurements of skin oscillations at selected locations (blue dots) elicited by an impulse (0.5 ms pulse width) applied at the digit III distal phalanx (DP) (red arrow). Right: PC spiking responses evoked by respective skin oscillations, shown for PC neuron model type 4. (C) Absolute Pearson correlation coefficients between skin oscillations shown in (B). (D) Magnitude spectra of skin oscillations shown in (B). (E) Reconstructed skin oscillations elicited by a bandpass-filtered noise stimulus (top trace, 50 to 800 Hz band) applied at the digit III DP. (F) Absolute Pearson correlation coefficients between skin oscillations at different distances from the contact location shown in (E). (G) Upper panel: PC mean firing rates elicited by an impulse applied at the digit III DP (red arrow; 15 µm max. peak-to-peak displacement across hand). Lower panel: cumulative percent of spikes (black) and responding PCs (blue) located within increasing distances from the contact location. Shaded region: results within digit III. (H) As in (G), for a 200 Hz sinusoidal stimulus (15 µm max. peak-to-peak displacement across hand). (I) As in (G), for a bandpass-filtered noise stimulus (50 to 800 Hz band, 5 µm max. RMS displacement across hand). (J) PC spiking responses (right, PC neuron model type 4) evoked by skin oscillations (middle) at selected locations (left, blue dots) elicited by a diharmonic stimulus (f_1_ = 50 Hz, f_2_ = 100 Hz) applied at the digit III DP (red arrow). Light blue bars: RMS displacements of skin oscillations; black and gray bars: percent of skin oscillation frequency magnitude spectrum composed of 50 Hz (black) or 100 Hz (gray) components; dark blue bars: PC firing rates calculated from spikes within shaded region. All plots show data from Participant 5 (P5).

As expected, due to the linearity of biomechanical transmission in the small signal regime^33,34^, we found the impulse measurements to accurately encode the transmission of skin oscillations in the hand (Figure S1). This allowed us to compute the whole-hand skin oscillations that would be evoked by an arbitrary input waveform by convolving the input waveform with the impulse measurements (see Methods; Figure S2A). Using this technique, we computed the skin oscillations evoked by diverse tactile input signals, including sinusoids, diharmonics, and bandpass-filtered noise. This method preserved the effects of biomechanical filtering, including the phase and amplitude of evoked skin oscillations (Figures 1E, 1F).

### Biomechanically mediated PC spiking activity

PC spiking responses are driven by deformations of the corpuscle caused by mechanical oscillations of surrounding tissues^31^. Thus, we sought to characterize the influence of biomechanical filtering on PC spiking responses. However, current experimental techniques preclude the *in vivo* measurement of PC population responses^32^. To overcome this limitation, we determined the spiking responses of whole-hand PC populations *in silico*, using the computed skin oscillations to drive a population of spiking PC neuron models that were fit to physiological data in prior research^35^ (Figure S2A). Each PC neuron model was driven by skin oscillations at its location in the hand. The spatial distribution of PCs across the hand was selected based on findings from a prior anatomical study^36^. We used this methodology to obtain whole-hand PC population spiking responses evoked by arbitrary tactile inputs supplied at any of four contact locations on the hand.

All stimuli evoked spiking activity in PCs throughout the hand, consistent with predictions from theory and findings from prior studies^3,6,11^. The majority of the elicited spiking activity and responding PCs were in hand regions far removed from the contact location. This was true for all stimulus types, including brief impulses (Figure 1G), sinusoids (Figure 1H and Figure S2C), and bandpass-filtered noise stimuli (Figure 1I and Figure S2E). In each case, the patterns of evoked spiking activity reflected the effects of biomechanical filtering (Figures S2B, S2D). For example, a brief impulse evoked PC spiking responses that varied based on hand location and exhibited sustained firing for more than 20 ms, reflecting the dispersive effects of biomechanical transmission (Figure 1B).

PC responses also exhibited characteristic entrainment behavior (phase-locking to the oscillations of periodic stimuli) that was modified by biomechanical filtering. For example, a diharmonic stimulus applied at the fingertip evoked distance-dependent patterns of entrainment (Figure 1J): Receptors near the contact location (< 60 mm) entrained to the higher frequency signal component (100 Hz), while more distant receptors entrained to the lower frequency component (50 Hz). This distance-dependent entrainment behavior arose due to the greater attenuation of higher frequency skin oscillations with distance (Figure 1J, black and gray bars), an effect of tissue viscoelasticity^2,26^. Thus, PC spiking activity is altered by effects of biomechanical filtering that vary with the location of the receptor in the hand.

### Location-specific biomechanical filtering

To more fully characterize the spatial dependence of biomechanical filtering, we analyzed the frequency-dependent transmission of skin oscillations across the hand. Applied sinusoidal stimuli (frequencies between 20 and 800 Hz) elicited widespread skin oscillations, with distance-dependent amplitudes that varied greatly with stimulus frequency (Figure 2A; results for other participants and contact locations: see Figure S3). At low frequencies (*≤* 100 Hz), the median transmission distance extended well beyond the stimulated digit, and the amplitude decay was non-monotonic with distance from the contact location. In contrast, higher-frequency components were concentrated within the stimulated digit and exhibited relatively monotonic decay with distance. We obtained similar findings for all participants and contact locations (Figure 2B). These complex, frequency-dependent patterns of biomechanical filtering are a function of soft tissue viscoelasticity and the heterogeneous morphology of the hand.

**Figure 2.**
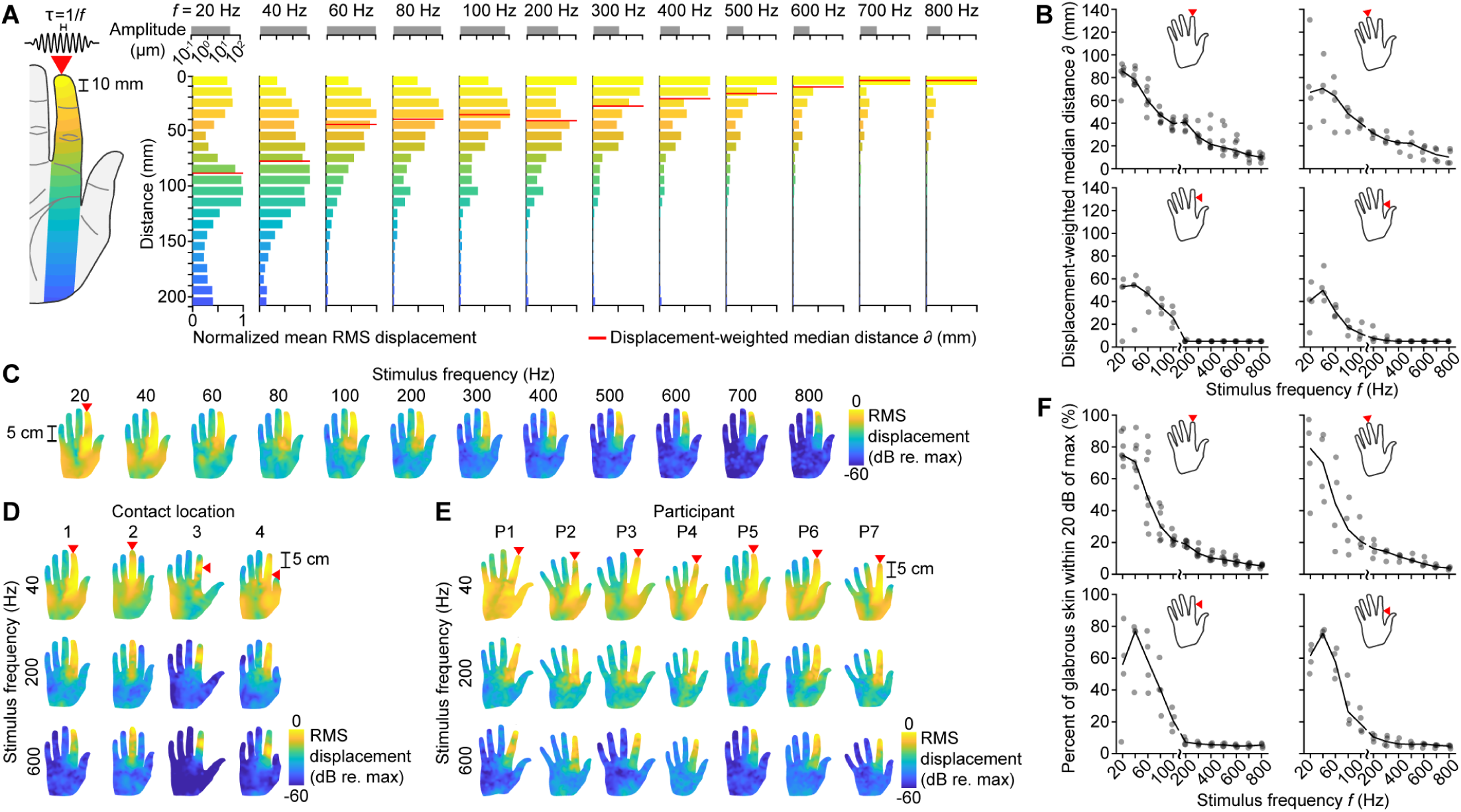
Biomechanical filtering in the hand is frequency- and location-dependent. (A) Normalized root mean square (RMS) skin displacements averaged within 10 mm-wide bands at increasing distances from the contact location elicited by sinusoidal stimuli of various frequencies (20 to 800 Hz). Amplitude scale bars (top, gray) show maximum peak-to-peak displacement across the hand at each frequency. Red lines: median transmission distance; red arrow: contact location. Shown for P5. (B) Median transmission distance of RMS skin displacement distributions across frequency for all participants and contact locations. Red arrows: contact location; lines: median across participants; dots: data points for each participant. (C) RMS displacement across the hand elicited by sinusoidal stimuli of various frequencies (20 to 800 Hz). Red arrow: contact location. Shown for P5. (D) As in (C), for other contact locations. Shown for P5. (E) As in (C), for P1 to P7. (F) Percent of glabrous skin where RMS skin displacement is within 20 dB of the maximum RMS displacement, for all participants and contact locations. Plots can be read as in (B).

We further observed the influence of the hand’s morphology and skeletal structure in the spatial patterns of biomechanical filtering across the whole hand (Figures 2C–2E; results for all participants and contact locations: see Figure S4). At low frequencies (*≤* 100 Hz), transmission was notably enhanced near the metacarpophalangeal (MCP) joint of the stimulated digit and in the lateral and contralateral extensions of the palmar surface (thenar and hypothenar eminences). These whole-hand patterns of biomechanical filtering also demonstrated that low-frequency stimuli evoked prominent oscillations (amplitudes within 20 dB of maximum) over a substantial fraction of the hand surface (mean 50 %; Figure 2F). In contrast, higher frequencies evoked skin oscillations that were confined to a smaller proportion of the hand surface (mean 8 %). These findings were consistent across contact locations and participants.

### Biomechanical filtering modulates PC tuning

We next examined how biomechanical filtering modulates the tuning characteristics of PCs, as reflected in their frequency-dependent entrainment behavior (Figure 1J). To do this, we analyzed whole-hand PC spiking activity evoked by sinusoidal stimuli with different frequencies (20 to 800 Hz). We characterized the frequency-dependent sensitivity of PCs by computing entrainment threshold curves, which represent the minimum displacement required to evoke entrainment at each frequency (see Methods). PCs located near the contact location exhibited U-shaped entrainment threshold curves with preferred (most sensitive) frequencies between 200 and 300 Hz (Figure 3A and Figure S5A). This result is consistent with established descriptions of PC function, which are based on *in vivo* experiments where the stimulus is applied adjacent to the PC^27–31^. However, we obtained diverging findings for the larger number of PCs located outside the contact region. Entrainment threshold curves for remote PCs varied greatly and exhibited multiple prominent minima, reflecting location-specific effects of biomechanical filtering (Figure 3A and Figures S5B, S5C). Moreover, PC entrainment threshold curves varied as the contact location varied (Figure 3B). Across the population, only a minority of PCs (≲ 20%) had preferred frequencies between 200 and 300 Hz.

**Figure 3.**
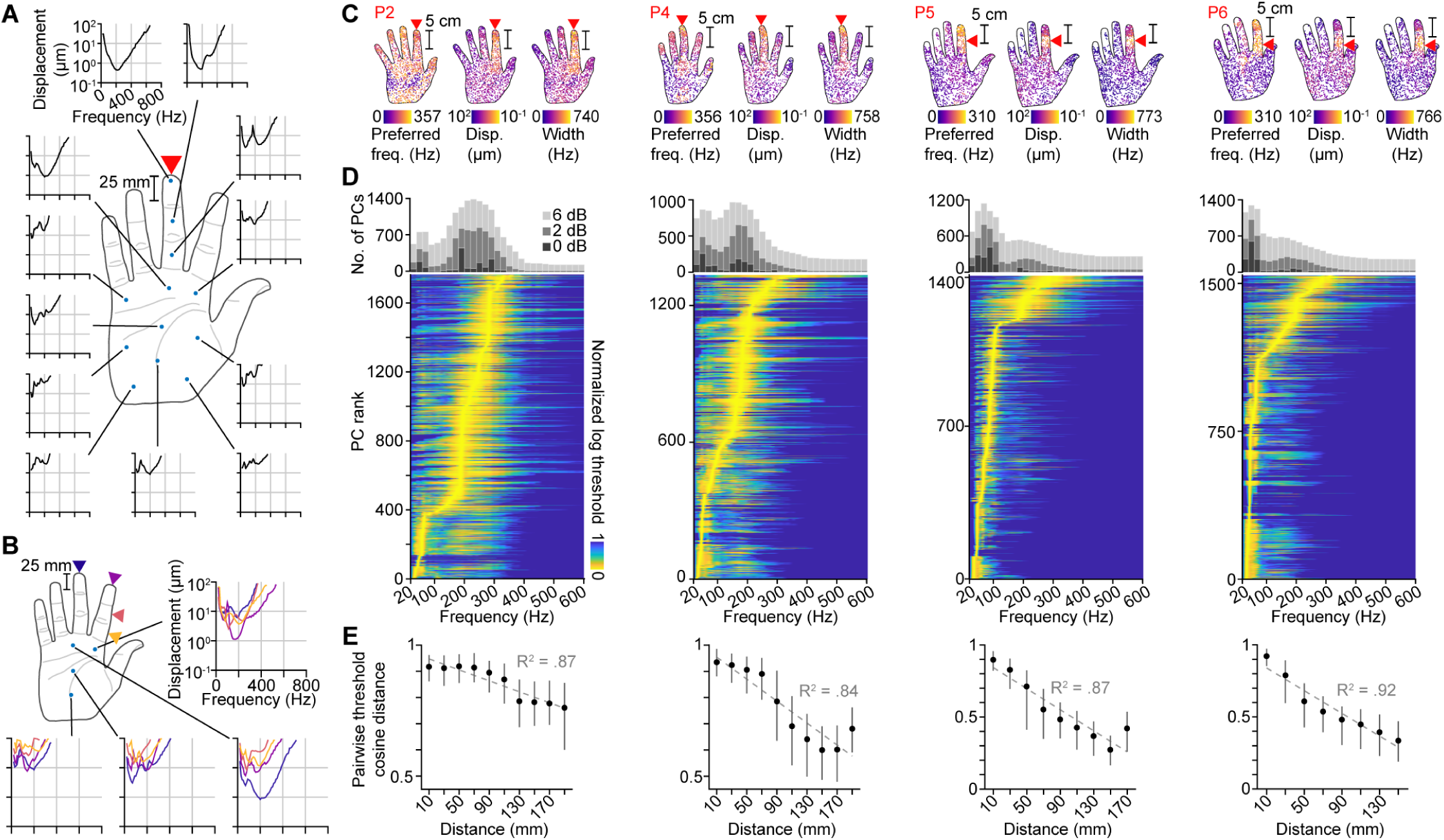
Biomechanical filtering diversifies PC response characteristics. (A) Entrainment threshold curves of PCs at selected locations (blue dots). Red arrow: Contact location. Shown for PC neuron model type 4 and P1. (B) Entrainment threshold curves of PCs at selected locations (blue dots) for each of four contact locations (colored arrows). Line colors correspond to contact locations. Shown for PC neuron model type 4 and P1. (C) Preferred frequency (left), minimum curve value (middle), and curve width (right) for each PC in the hand. Red arrow: contact location; red text: participant number. (D) Lower panel: Entrainment threshold curves for all PCs in the hand rank ordered by preferred frequency. Upper panel: Number of PCs at each frequency with entrainment threshold curve values within 0 dB (light gray), +2 dB (medium gray), and +6 dB (dark gray) of the curve minimum. Participants and contact locations as in (C). (E) Cosine distances between all pairs of entrainment threshold curves of PCs located within 10 mm of the contact location and those of PCs located within 20 mm-wide bands at increasing distances from the contact location. Center distance for each band shown. Dots: median; error bars: IQR; gray dotted lines: linear fit of medians; gray text: R^2^ value for linear fit. Participants and contact locations as in (C).

We further analyzed the distribution of frequency tuning across whole-hand PC populations by rank ordering all entrainment threshold curves by preferred frequency (Figures 3C, 3D; results for all participants and contact locations: see Figure S6). Within each population, PCs exhibited diverse preferred frequencies, ranging from 25 to 420 Hz (Figures S7A–S7D). The preferred frequencies of PCs located near the contact location were consistent with values obtained in prior studies (200 to 300 Hz), as noted above. However, PCs located outside the contact region exhibited a wider range of frequency sensitivities (Figures S7E–S7H). Strikingly, across all participants and contact locations, a substantial proportion of PCs in a population preferred frequencies below 100 Hz (mean 47 %). In addition, PCs at greater distances from the contact location exhibited more narrowly tuned curves and higher thresholds, indicating greater specificity in frequency preference and lower sensitivity (Figure 3C).

However, the entrainment threshold curves exhibited complex shapes that varied greatly with hand location in a manner not adequately summarized by preferred frequency, curve width, or minimum threshold. To characterize distance-dependent variations in the threshold curves, we instead calculated the pairwise cosine distance between threshold curves of PCs at the contact location and those within regions at progressively greater distances from the contact location (Figure 3E; results for all participants and contact locations: Figure S8). For all participants and contact locations, the median cosine distance between threshold curves decreased with increasing distance from the contact location (0.021 to 0.095 per 20 mm; R^2^ = 0.51 to 0.95). Together, these findings demonstrate that pre-neuronal biomechanical filtering diversifies frequency response characteristics in whole-hand PC populations.

### Biomechanical filtering diversifies tactile encoding

We next asked whether this diversification enhanced information encoding in PC population spiking responses, particularly for the large proportion of PCs outside the contact region. To address this, we characterized the dimensionality and information content of PC population spiking activity as a function of distance from the contact location. Informed by prior research^37,38^, we employed a diverse set of stimuli containing sinusoidal, diharmonic, and bandpass-filtered noise signals with various frequency and amplitude parameters (Tables S1–S3). We used principal component analysis to assess the latent dimensionality of PC firing rates in subpopulations of PCs within increasing maximum distances from the contact location. For all participants and contact locations, dimensionality, calculated as the number of principal components needed to capture 99 % of the variance in the firing rates, increased with distance. The dimensionality was 2 to 5 times higher at distances greater than 100 mm from the contact location than at distances less than 20 mm (Figure 4A). Thus, PCs at increasing distances from the contact location captured progressively more variance, highlighting the facilitative role of biomechanical filtering in PC population encoding.

**Figure 4.**
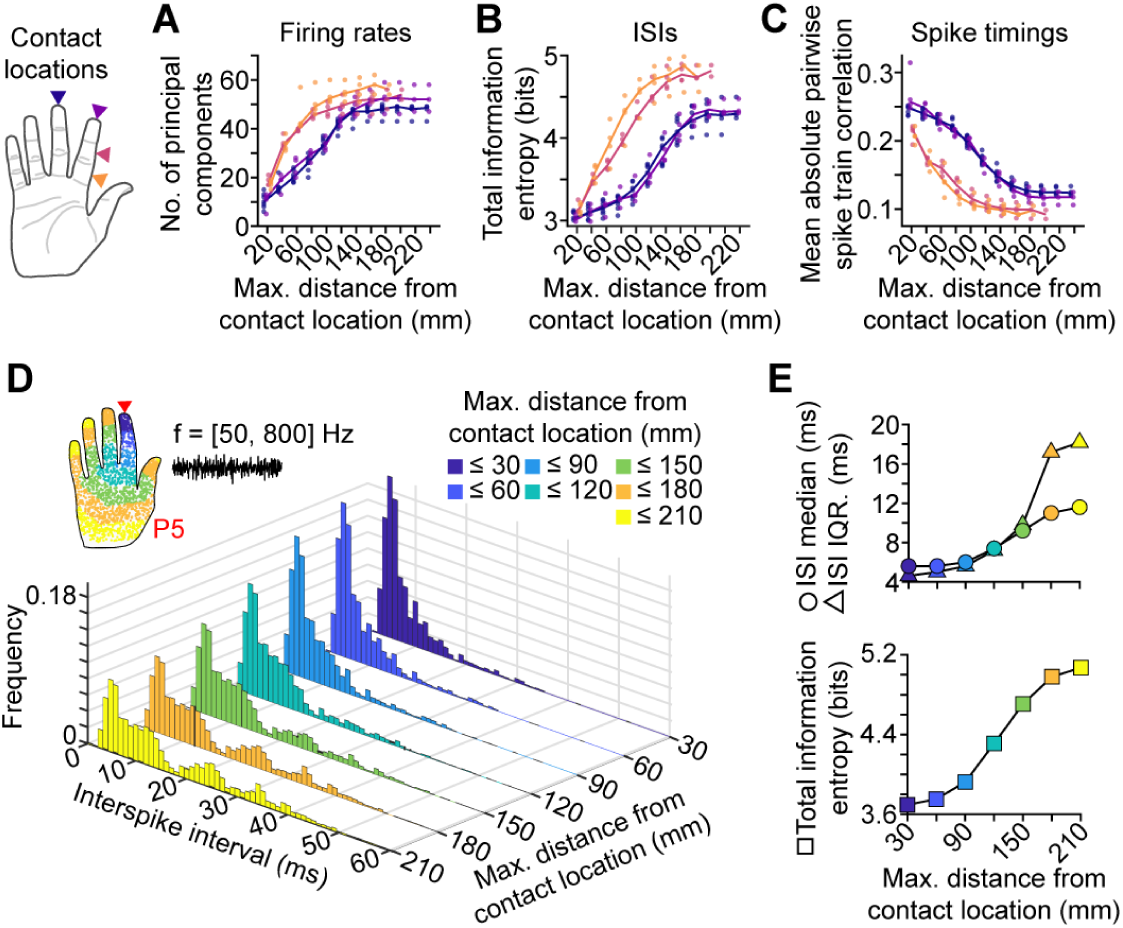
Biomechanical filtering diversifies PC spiking activity. (A) Number of principal components explaining 99 % of the variance in the firing rates of PCs within increasing distance ranges from the contact location. Colors indicate contact location; dots: data points for each participant; lines: median across participants. Analyses are conducted on PC spiking activity evoked by a diverse stimulus set. (B) Total information entropy of interspike interval (ISI) distributions (1 ms bin width) constructed from the spiking activity of PCs within increasing distance ranges from the contact location. Plot can be read as in (A). (C) Mean absolute spike train correlation between all pairs of PCs both located within increasing distances from the contact location. Spike trains were binned with a bin width of 1 ms. Plot can be read as in (A). (D) ISI histograms constructed from the spiking activity of PCs located within increasing distances from the contact location (hand inset) in response to a bandpass-filtered noise stimulus (50 to 800 Hz band, 5 µm max. RMS displacement across hand, 175 ms duration) applied at the digit II DP of P5. (E) Median (circles), interquartile range (triangles), and total information entropy (squares) of the ISI histograms in (D).

In addition to firing rates, PCs signal information about touch events via spike timing^38^. We characterized information encoded in PC spike timing by computing the Shannon information entropy of interspike interval (ISI) histograms constructed from the spiking activity of PC subpopulations within increasing maximum distances from the contact location. For all stimuli, ISIs were generally larger and more broadly distributed with increasing distance (Figures 4D, 4E and Figure S9). Consequently, information encoded in the ISIs increased monotonically with distance by a factor of more than 1.4 before plateauing at 140 to 180 mm from the contact location (Figure 4B). The findings were robust to variations in ISI histogram bin widths (Figures S10A–S10G).

We also analyzed the correlations between spiking responses evoked in subpopulations of PCs within increasing maximum distances from the contact location. Consistent with the ISI findings, as distance increased, the spiking responses of remote PCs became progressively less correlated with the spiking responses of PCs near the contact location, as quantified by mean pairwise spike train correlations (Figure 4C and Figures S10H–S10N). Together, these information encoding measures demonstrate that biomechanical filtering diversifies PC spiking activity by enabling remote PCs to encode information not captured by PCs near the contact location, thereby supporting tactile encoding efficiency^39,40^. The plateaus in the computed measures also indicate that some response redundancy is preserved within the population responses.

## DISCUSSION

Our study combined high-resolution vibrometry measurements of whole-hand biomechanical transmission with neural simulations leveraging extensively validated neuron models^35^. The results shed light on the pre-neuronal role of biomechanical filtering in modulating and diversifying PC population spiking activity. These findings indicate that previously documented response characteristics of isolated PCs, including entrainment threshold curves, do not accurately capture the behavior of most PCs within the human hand. Because the majority of PCs that respond during touch events are distant from the contact location, neural activity in remote PCs represents a dominant proportion of the total population response and can thus be expected to affect downstream tactile processing and, ultimately, perception.

The perceptual significance of widespread PC responses is well established by studies that reveal prominent vibrotactile summation and masking effects arising between stimuli applied at distant hand locations^18,19^. There is also ample evidence that biomechanically facilitated activity in remote PCs influences perception. Prior experiments have demonstrated that in subjects with impaired tactile sensation due to anesthesia, nerve compression, or traumatic injury, tactile perception in both the high-frequency (> 80 Hz) and flutter range (*∼* 20 Hz) can be mediated by remote PCs^17,41,42^. Prior research also shows that fine surface textures explored with the anesthetized finger can be perceptually discriminated^16^. These discriminative abilities were found to be facilitated by spiking activity in remote PCs evoked by skin oscillations transmitted far from the contact location^6^.

Prior studies have also demonstrated the significance of biomechanical transmission for the perception of artificial haptic feedback. For example, the propensity of lower-frequency skin oscillations to travel greater distances has been exploited to realize frequency-controlled haptic effects that are perceived to expand or contract in spatial extent from a single localized site of stimulation^26^. Moreover, a recent study reported prominent effects of biomechanical transmission on the perception of tactile motion stimuli supplied via airborne ultrasound^43^.

Accounting for the effects of biomechanics on peripheral tactile encoding may also shed light on touch perception in other settings. For example, perceived vibrotactile intensity is highly frequency-dependent^44,45^, but the observed dependence conflicts with predictions derived based on responses of PCs adjacent to the contact location^37^. Moreover, our understanding of the perception of polyharmonic stimuli is incomplete. A proposed model for the perception of stimuli with complex frequency spectra invoked the existence of mechanoreceptor subpopulations that vary in frequency tuning^46,47^. However, this model has not gained traction because such subpopulations have not been previously observed. Indeed, established characterizations of PC function based on stimulation near the receptor location depict PC frequency sensitivity as highly stereotyped.

Our findings may also shed light on a number of peculiar aspects of PC innervation of the hand. Despite the stereotyped response properties of isolated PCs and their large receptive fields, which span most of the hand, PCs in the glabrous skin number in the hundreds or more^36,48–50^. Together, these characteristics may be interpreted to imply tremendous response redundancy, which would be at odds with encoding efficiency hypotheses^39,40^. However, our results demonstrate that biomechanical filtering diversifies PC response characteristics, thereby reducing PC population response redundancy and enhancing encoding efficiency. Furthermore, prominent clusters of PCs are observed near the MCP joints in human hands^49,50^. Near those locations, we observed elevated oscillation amplitudes at low frequencies (< 100 Hz), suggesting that PCs may be preferentially located in regions of the hand where biomechanical transmission is facilitated.

The frequency-dependent patterns of biomechanical transmission and filtering we observed are generally consistent with prior characterizations of mechanical propagation near the contact location^11,26,51^, taking into account likely differences in contact conditions. Here, we present whole-hand measurements at significantly greater spatiotemporal resolution than was used in prior studies. This made it possible to resolve previously unobserved effects of biomechanical transmission and filtering throughout the hand, including the non-monotonic decay of oscillation amplitude with distance and the contact location-dependent variations in filtering.

Despite the observed complexity of biomechanical transmission in the hand, several characteristics were conserved across multiple hands and contact locations. These included the frequency-dependent variations in transmission amplitude with distance, the elevated transmission distances at low frequencies (*≤* 100 Hz), and the enhancement of transmission near the MCP joints. These features demonstrate how biomechanical filtering generates spatial and spectral structure that the brain could learn and exploit, similar to hypotheses for efficient encoding of whole-hand touch events^52^, object slippage^53^, and tool use^8^.

Overall, the pronounced effects of biomechanics that we observed in the PC system exemplify how pre-neuronal mechanisms can play a crucial role in sensory processing, which is supported by analogous findings in other sensory systems. For example, the effect of biomechanical filtering on tuning properties across PC populations in the hand is somewhat comparable to the frequency-place transform that is biomechanically affected by the mammalian cochlea^54,55^. Moreover, the biomechanics of the human body filters the frequency content of motion inputs to the vestibular system during natural movement, which likely has consequences for underlying neural circuitry^56^. Similar conclusions about the impact of biomechanics in sensory processing have been drawn in studies of the rodent vibrissal system, where the mechanics of the whiskers are instrumental to tactile neural coding^57,58^.

## METHODS

### *In vivo* optical vibrometry

Mechanical oscillations across the volar hand surface were imaged with a scanning laser Doppler vibrometer (SLDV; model PSV-500, Polytec, Inc., Irvine, CA; sample frequency 20 kHz) fastened to a pneumatically isolated table. During each recording, the hand was fixed to the table in an open, palm-up posture via custom-fit 3D-printed supports that were fastened to the table and adhered to the fingernails of all but the stimulated digit (Figure 1A). Participants (n = 7, 5 male) were 20 to 45 years of age (mean 27.4 years) and were recruited from the student and staff population at the authors’ institution. The sample size was determined based on previously published research employing similar methodologies^10,11,26,52^. Participants were seated in a reclined chair with the arm relaxed, supported by a foam armrest, and constrained with Velcro straps. All participants gave their informed, written consent prior to the data collection. The study was approved by the Human Subjects Committee of the University of California, Santa Barbara (Protocol Number 9-18-0676).

The SLDV imaged spatially and temporally resolved skin oscillations at sampling locations distributed on a uniform grid covering the entire volar hand surface (grid spacing 8 mm, 200 to 350 locations). The sampling grid exceeded the Nyquist criterion threshold for frequencies in the tactile range (0 to 1000 Hz), at which spatial wavelengths are between 20 to 100 mm^52^. Oscillations were imaged in the normal direction to the skin surface. Prior vibrometry measurements have demonstrated that most of the energy in evoked skin oscillations is concentrated in oscillations normal to the skin surface^26^ and that stress in the normal direction is highly predictive of PC spiking responses^35^.

All data were captured from the right hands of participants. Hand lengths ranged from 18 to 21.6 cm (mean 19.9 cm) as measured from the tip of digit III to the bottom of the hand at the middle of the wrist. Each hand was positioned 36 cm below the SLDV aperture, which ensured that the measurements captured at least 95 % of the signal variance at all measurement locations. Each participant’s hand shape and the 2D spatial coordinates of all measurement locations were captured via the integrated SLDV geometry processor and camera. Measurements were interpolated to obtain skin oscillations at other locations on the 2D hand surface (see Supplemental Text).

Measured skin oscillations were evoked by mechanical impulses (rectangular pulse, duration 0.5 ms) applied at each of the four contact locations described below. Measurements were synchronized to the stimulus onset. Each measurement was obtained as the median of 10 captures and bandpass filtered to the vibrotactile frequency range (20 to 1000 Hz). Frequency analysis was performed by computing magnitude spectra, which were smoothed using a moving median window (width: 3 samples) to remove measurement artifacts. Numerical frequency-domain integration was employed to obtain skin displacement from velocity. Stimuli were delivered via an electrodynamic actuator (Type 4810, Brüel & Kjær) driven with a laboratory amplifier (PA-138, Labworks). The actuator terminated in a plastic probe (7 *×* 7 mm contact surface) that was adhesively attached to the skin at the stimulus contact location. The actuator and probe were configured to avoid obstructing the optical path used for the SLDV measurements.

Stimuli were applied at each of four different contact locations (CL) that were registered to standard anatomical positions on the hand: the distal phalanx (DP) of digit II along the axis of the finger (CL 1, n = 7 participants), the DP of digit III along the axis of the finger (CL 2, n = 4), the intermediate phalanx (IP) of digit II perpendicular to the axis of the finger (CL 3, n = 4), and the proximal phalanx (PP) of digit II perpendicular to the axis of the finger (CL 4, n = 4) (Figure 1A). These measurements took approximately 10 minutes per contact condition per participant. Measurements for CL 2, 3, and 4 were captured from a subset of participants from which measurements for CL 1 were captured (P1, P4, P5, and P6).

### Computing skin oscillations evoked by arbitrary stimuli

Theory and experimental findings^33,34^ indicate that biomechanical transmission in the hand is linear for the stimulus magnitudes employed here. Consequently, the transmission of evoked skin oscillations may be mathematically described by a wave equation of the form L u(**x**, t) = 0, where L is a linear partial differential operator encoding wave transport, **x** is a skin location, t is time, and u(**x**, t) is the evoked skin oscillation pattern. From linear systems theory, an arbitrary force stimulus F(t) applied to the skin at location **x**_0_ evokes oscillations u(**x**, t) given by

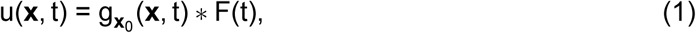

where *∗* denotes convolution in time and g**_x_**_0_ (**x**, t) is the Green’s function encoding the excitation of skin oscillations evoked by a unit Dirac impulse applied at **x**_0_. We empirically determined the Green’s functions for each hand and contact location **x**_0_ using the impulse-driven skin oscillation measurements described above. The skin oscillations evoked by arbitrary stimuli F(t) were numerically computed using Equation 1.

To confirm the accuracy of this method, we compared the numerically computed oscillations with measured oscillations evoked by sinusoidal stimuli F(t) with frequencies ranging from 20 to 640 Hz. The measurement procedure was identical to the one described above, apart from the input waveform. Consistent with linear systems theory, we found that the numerically computed oscillations closely approximated the actual measurements at all frequencies (Figure S1). Because the numerical methodology avoids the need for time-intensive experiments, we used it to determine skin oscillations evoked by arbitrary stimuli in the remainder of the experiments.

### Stimuli

We analyzed skin oscillations u(**x**, t) evoked by sinusoidal, diharmonic, and bandpass-filtered noise stimulus waveforms F(t). For sinusoidal stimuli, F(t) = F_0_ sin (2*π*ft), where f is frequency and F_0_ is force amplitude. Force amplitudes were approximately 0.25 N, except as noted. For diharmonic stimuli, F(t) = F_1_sin(2*π*f_1_t) + F_2_sin(2*π*f_2_t), with independent force amplitudes F_1_ and F_2_. F_1_ and F_2_ were selected to ensure that the maximum evoked peak-to-peak skin displacements D_pp_ were equal, where

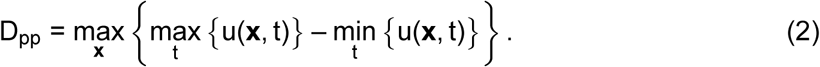

Bandpass-filtered noise stimuli were synthesized using a spectral Gaussian white noise algorithm^59^, then bandpass filtered to the desired frequency range. Each bandpass-filtered noise stimulus was generated from the same Gaussian white noise trace, then scaled by a force amplitude F_0_.

### Whole-hand neural simulations

PC population spiking responses were obtained *in silico* by using the skin oscillations to drive a population of spiking neuron models extracted from a simulation package (Touchsim^35^, Python) (Figure S2A), similar to the methodology applied in prior work investigating PC population responses during whole-hand touch events^13^. The PC neuron models consisted of four PC types, each trained and validated in a prior study on electrophysiology recordings from non-human primates^37^. Each PC neuron model type varies slightly in its response properties (Figure S5A). The PC neuron models supply a nonlinear mapping from skin displacement to spiking output and accurately reproduce experimentally identified PC response characteristics, including response thresholds that vary across several orders of magnitude over the vibrotactile frequency range^28,30^ and frequency-dependent thresholds of entrainment^27,29,31^. The stimulus amplitudes used in this work fell within the range over which the PC models were validated.

Whole-hand PC populations were assembled by sampling a random distribution weighted by spatial densities *σ* reported in prior studies^36,48^: *σ* = 25 cm^2^ in the distal phalanges and *σ* = 10 cm^2^ in the rest of the hand. Except where otherwise noted, the PC neuron model type for each PC in a population was randomly selected to be one of the four PC neuron model types noted above. Each PC was driven by the time-varying skin oscillations u(**x**_m_, t), where **x**_m_ is the PC location. This produced a spike train represented as an ordered sequence Y_m_ = {t_1_, t_2_, …, t_Q_}, where t_i_ are spike times and Q is the number of stimulus-evoked spikes.

### PC entrainment threshold curves

Entrainment threshold curves were constructed to characterize PC frequency sensitivity. At each frequency (20 to 800 Hz), the force amplitude of the sinusoidal input stimulus was varied to determine the entrainment threshold for each PC in the whole-hand population. The entrainment threshold was identified as the minimum input force amplitude at which one spike was elicited per cycle of the sinusoidal stimulus. For each mth PC and frequency f, the threshold curve E_m_(f) recorded the maximum peak-to-peak displacement of skin oscillations evoked across the hand (D_pp_, Eq. 2) at the identified entrainment force amplitude. The maximum D_pp_ across all conditions was 100 µm. In prior literature, threshold curves were determined by placing the stimulating probe near the hotspot of the terminating neuron^27–31^ (Figure S5A). Here, however, we analyzed threshold curves for all PCs across the hand with the stimulus contact location held constant. This preserved the effects of biomechanical transmission and filtering that were not captured in prior approaches. For each PC, the preferred (most sensitive) frequency was computed as f*^∗^* = arg min_f_{E_m_(f)}. The width of each threshold curve was determined as the full width of the threshold curve (not necessarily contiguous) at half-minimum and characterized the sensitivity bandwidth of the respective PC.

### Cosine distance analysis

The pairwise similarity of entrainment threshold curves for different PCs was assessed using the cosine distance c_ij_,

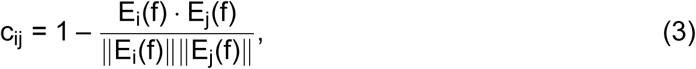

where i and j index the threshold curves of PCs in a population. We conducted this analysis by separating whole-hand PC populations into PC subpopulations P_k_ located within different distance ranges from the stimulus contact location, where k = 0, 1, *. . .* , K indexes the distance range. P_0_ contained all N_0_ PCs located within 10 mm of the contact location. For k > 0, P_k_ contained all N_k_ PCs located at distances d_k_ from the contact location satisfying (20(k–1)+10) *≤* d_k_ < (20k+10) mm. Distances on the 2D hand surface were computed from the SLDV geometry data using Djikstra’s algorithm. We computed the mean cosine distance *σ*_k_ between all PC threshold curves in P_0_ and all PC threshold curves in P_k_ as

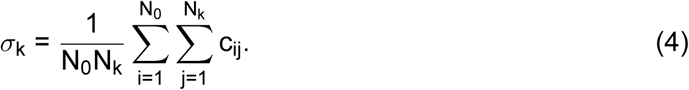

We computed the mean pairwise cosine distance *σ*_0_ between PC threshold curves in P_0_ as

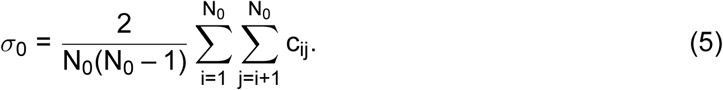

### PC population spiking activity analysis

Efficient encoding hypotheses posit that neural sensory circuitry should minimize redundancy^39,40^. We assessed encoding efficiency by analyzing mean firing rates and spike timing, both of which are salient to tactile encoding^60–62^. To this end, we assembled a diverse set of stimuli encompassing commonly occurring tactile signals based on prior studies^37,38^. The stimulus set contained 60 sinusoidal, 117 diharmonic, and 50 bandpass-filtered noise stimuli that varied in amplitude and frequency parameters (see Supplemental Text and Tables S1–S3). We obtained spike trains evoked in whole-hand PC populations for each stimulus (n = 227), contact location (n = 4), and participant (n = 4 or n = 7 depending on the contact location).

To assess the redundancy in spiking responses of remotely located PCs, we constructed PC subpopulations P^r^ located within increasing distances d_r_ = 20r mm from the contact location. The subpopulations formed a nested array of sets, P^1^ *⊂* P^2^ *⊂ · · · ⊂* P^R^, successively encompassing larger numbers of PCs, N^r^, where N^r^ = N^r–1^ + ΔN^r^. We analyzed spiking responses from each subpopulation using principal component analysis (PCA), interspike interval (ISI) information entropy, and spike train correlations.

### Firing rate latent dimensionality analysis

To assess the latent dimensionality of spiking activity from each subpopulation r, principal component analysis (PCA) was performed on the matrix of time-averaged firing rates 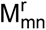, where m indexed PCs and n indexed the stimuli. The data matrix 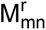 was standardized across stimuli (zero-mean and unit standard deviation). To assess the latent dimensionality of the subpopulation firing rates, we computed the minimum number B(r) of principal components that captured at least 99 % of the variance. A higher value of B(r) indicated greater firing rate heterogeneity in subpopulation r.

### Interspike interval information entropy analysis

We computed interspike intervals (ISIs) from spike trains 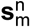 evoked in each PC m by each stimulus n. From these data, we computed normalized ISI histograms for each PC subpopulation P^r^. In the main results (Figure 4B), the histogram bin size was Δt = 1 ms because PCs encode touch information with millisecond precision^38^. We obtained similar results for values of Δt between 0.5 and 2 ms (Figures S10A–S10G). Let 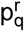 be the probability of an ISI t from P^r^ falling in the range t_q_ *≤* t < t_q+1_, where t_q_ = q Δt. We assessed information in spike timing by computing the Shannon information (entropy) H(r), given by

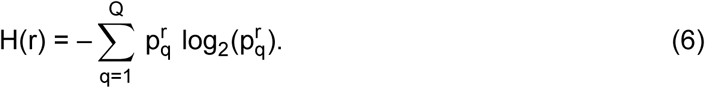

We also applied the same analysis to individual stimuli (Figure 4D and Figure S9). Higher ISI entropy values H(r) indicated more information and less redundancy in spike timing activity.

### Spike train correlation analysis

We computed spike train correlations^63–65^ using binned spike train vectors with bin width Δt = 1 ms (Python package elephant^66^) (Figure 4C). We obtained similar results for values of Δt between 0.5 and 2 ms (Figures S10H–S10N). For each stimulus n and PC subpopulation P^r^, we assessed pairwise spike train correlations by computing Pearson correlation coefficients 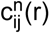 between each pair of binned spike trains 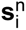 and 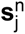, where i and j index PCs in P^r^. We computed the mean spike train correlation c(r) for each subpopulation P^r^ using

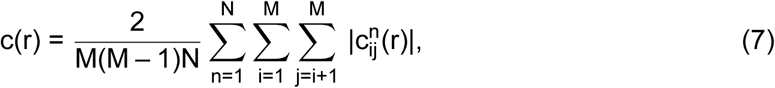

where M is the number of PCs in a subpopulation P^r^ and N is the number of stimuli. Lower spike train correlations indicated less redundancy within population spike timing activity. In contrast to the ISI entropy analysis, spike train correlations took into account the relative differences in spike times between different PCs.

## Data and Code Availability

The data and code that support the findings of this study are available on Zenodo: https://doi.org/10.5281/zenodo.10637210.

## Acknowledgements

We thank Benoit Delhaye for feedback and Polytec Inc. for the use of a vibrometry system. This research was supported by National Science Foundation award 1751348 to Y.V. and Leverhulme Trust research project grant RPG-2022-031 to H.P.S. N.T. was supported by a Link Foundation Modeling, Simulation, and Training Fellowship and a UC Santa Barbara Graduate Opportunity Fellowship.

## Author Contributions

N.T., G.R., H.P.S., Y.V., conceptualization, visualization, writing – original draft, and writing – review & editing; N.T. and G.R., data curation and validation; N.T., formal analysis; Y.V., funding acquisition; B.D., Y.S., and Y.V., investigation; N.T., G.R., B.D., Y.S., H.P.S., and Y.V., methodology; N.T., G.R., and B.D., software; H.P.S. and Y.V., supervision.

## Supplemental Text

### Skin oscillation interpolation across hand surface

Green’s functions were measured at discrete locations **x**_k_ in 2D pixel space and mapped to real space using participants’ hand lengths. The sampling grid exceeded the Nyquist criterion threshold for frequencies in the tactile range (0 to 1000 Hz), which exhibit spatial wavelengths between 20 to 100 mm^52^. This enabled us to determine Green’s functions and skin oscillations at arbitrary locations **x** via interpolation. Data was first extrapolated to the boundary of the 2D hand surface outside the convex hull bounded by the measurement locations using distance weighting. The skin displacements at boundary locations **x**_b_ were computed as

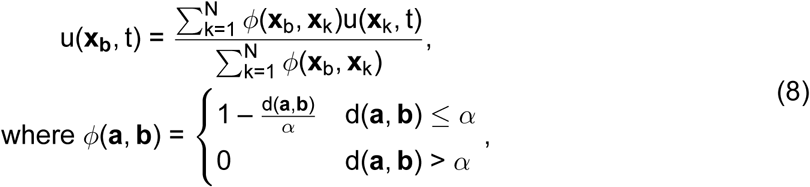

where d(**a**, **b**) was the Euclidean distance between locations **a** and **b** in pixel space and N was the total number of measured locations. We selected *α* = 90 pixels to ensure at least two measured locations contributed to the extrapolation of the displacement at boundary locations. The extrapolated displacements u(**x**_b_, t) and the sampled measurements u(**x**_k_, t) were used to compute displacements at arbitrary locations u(**x**, t) inside the 2D hand surface using nearest neighbor interpolation.

### Stimulus set for PC population spiking activity analysis

The stimulus set consisted of 60 sinusoidal, 117 diharmonic, and 50 bandpass-filtered noise stimuli presented at various amplitudes. The sinusoidal stimuli were 100 ms in duration and comprised 12 frequencies and 5 amplitudes per frequency (stimulus parameters: see Table S1). The diharmonic stimuli were 100 ms in duration and comprised 13 frequency pairs and 9 amplitude combinations per pair (Table S2). The bandpass-filtered noise stimuli were 1000 ms in duration and comprised 10 frequency bands and 5 amplitudes per band (Table S3).

For the sinusoidal stimuli, the force amplitudes of the stimuli were selected to yield skin oscillations with a specific maximum peak-to-peak displacement across all hand locations (D_pp_, Eq. 2). For the diharmonic stimuli, force amplitudes were selected independently for each sinusoidal component to yield specified values of D^1^ and D^2^ , the maximum peak-to-peak displacement across all hand locations for each sinusoidal component separately (see Methods). For the bandpass-filtered noise stimuli, we instead determined the force amplitude by specifying D_RMS_, the maximum RMS displacement of skin oscillations across all hand locations. The maximum peak-to-peak displacement across the whole hand evoked by any stimulus in the stimulus set was 200 µm.

For the sinusoidal and diharmonic stimuli, the minimum D_pp_ for a given frequency component was the smallest displacement that elicited entrainment in one of the four PC models via direct stimulation (i.e., without the effects of biomechanical filtering). The exception was 800 Hz for which D_pp_ = 10 µm, which was below the entrainment threshold at that frequency. The maximum D_pp_ for each frequency component was either 100 µm or the D_pp_ which yielded a maximum peak-to-peak skin acceleration of 50 *×* g = 490 m/s^2^, whichever was smaller. This ensured that displacements remained in a regime where the skin could be considered approximately linear and in which the PC models were validated. For sinusoidal stimuli, the 5 amplitudes were equally spaced in log space between the identified minimum and maximum D_pp_. For diharmonic stimuli, 3 amplitudes were selected for each frequency component and equally spaced in log space between the identified minimum and maximum D_pp_. Stimuli were then presented for each combination of amplitudes (3 *×* 3 = 9 amplitude combinations per frequency pair). For each bandpass-filtered noise stimulus, the selected D_RMS_ were 0.5, 1, 5, 10, and 20 µm.

## Supplemental Figures

**Figure S1.**
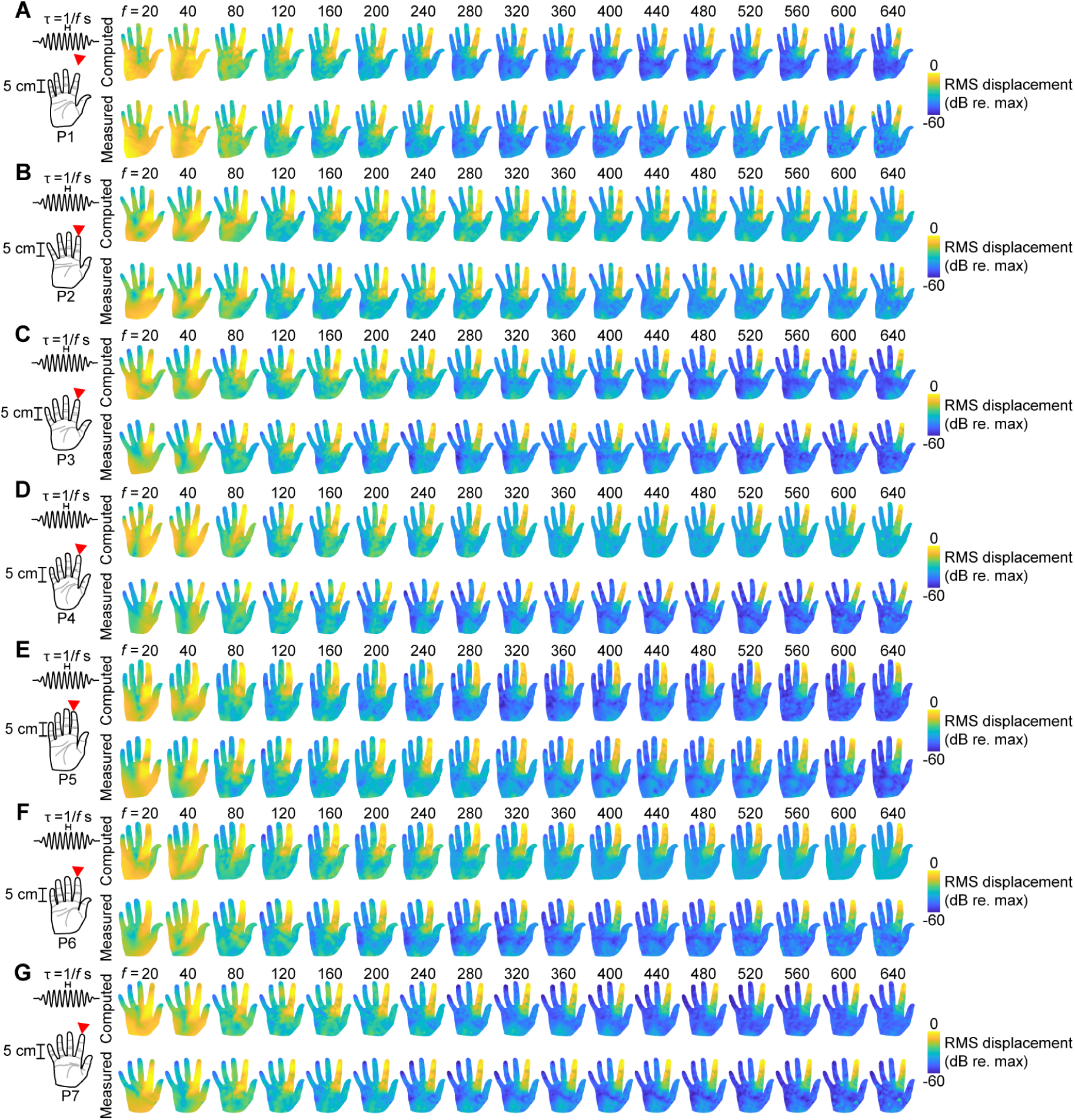
Numerically determined versus measured whole-hand RMS skin displacements across frequency. (A) Normalized root mean square (RMS) whole-hand skin displacements (log scale) elicited by windowed sinusoidal stimuli of various frequencies (20 to 640 Hz) applied at the digit II distal phalanx (DP) of Participant 1 (P1). Top row: numerically determined skin oscillations (Materials and Methods). Bottom row: experimentally measured skin oscillations. (B) As in (A) for P2. (C) As in (A) for P3. (D) As in (A) for P4. (E) As in (A) for P5. (F) As in (A) for P6. (G) As in (A) for P7.

**Figure S2.**
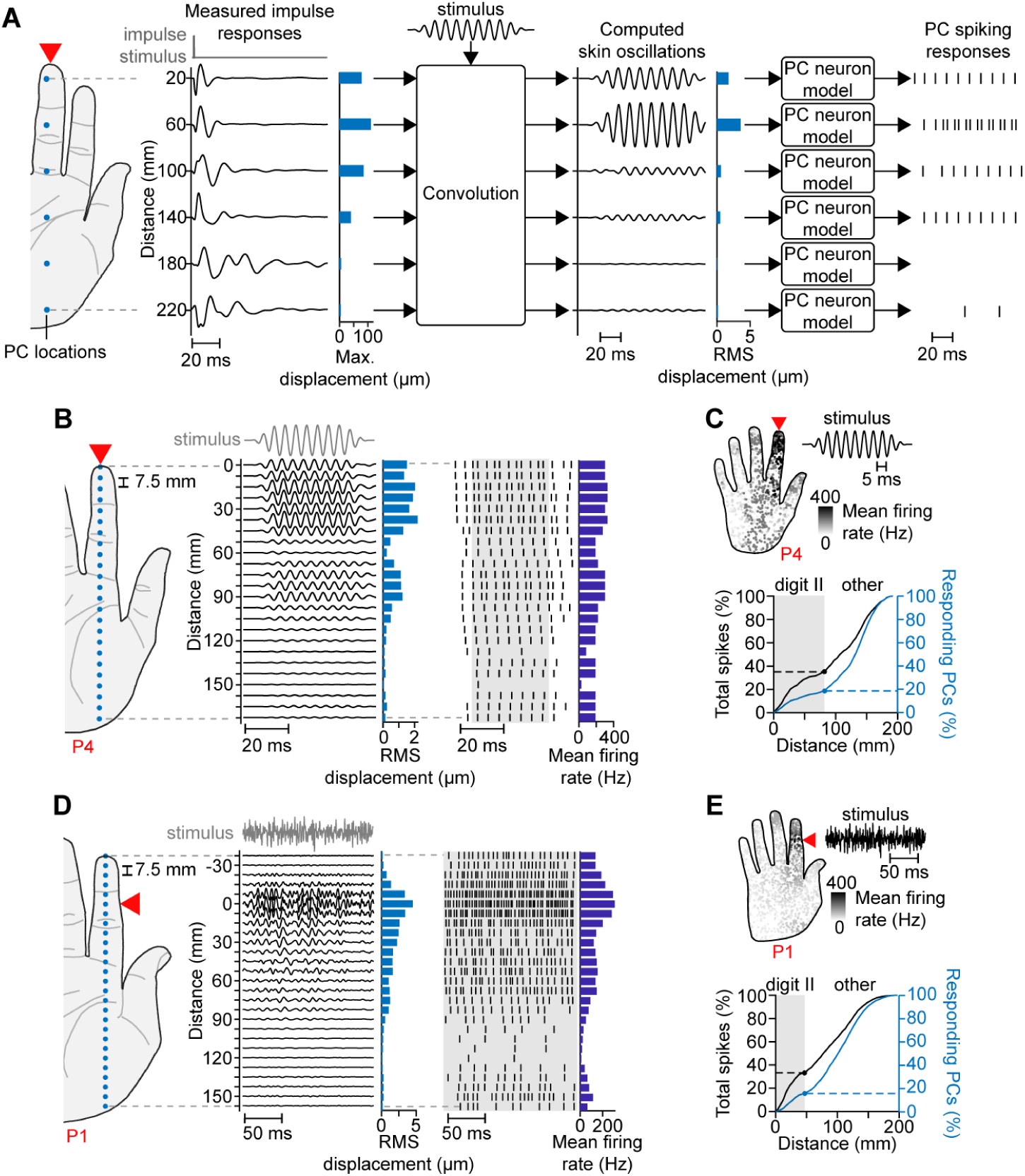
Measurement-driven neural simulation methodology. (A) Skin oscillations are computed at PC locations (blue dots) in response to an arbitrary stimulus via convolution with measured impulse responses (Materials and Methods). The computed skin oscillations are input to integrate-and-fire PC neuron models to produce spiking responses. Here, the stimulus is a 100 Hz sinusoid applied at the digit III DP of P5. (B) PC spiking responses (right) evoked by skin oscillations (middle) at selected locations (left, blue dots) elicited by a 100 Hz sinusoidal stimulus applied at the digit III DP of P4 (10 µm max. peak-to-peak displacement across hand). Shown for PC model type 4. Light blue bars: RMS skin displacements; dark blue bars: PC mean firing rates calculated from spikes within the shaded region. (C) Upper panel: PC mean firing rates elicited by the stimulus in (B). Lower panel: cumulative percent of total spikes (black) and responding PCs (blue) located within increasing distances from the contact location. Shaded region: results within digit II. (D) As in (B) for a bandpass noise stimulus (20 to 800 Hz band, 5 µm max. RMS displacement across hand) applied at the digit II intermediate phalanx (IP) of P1. (E) As in (C) for the stimulus in (D).

**Figure S3.**
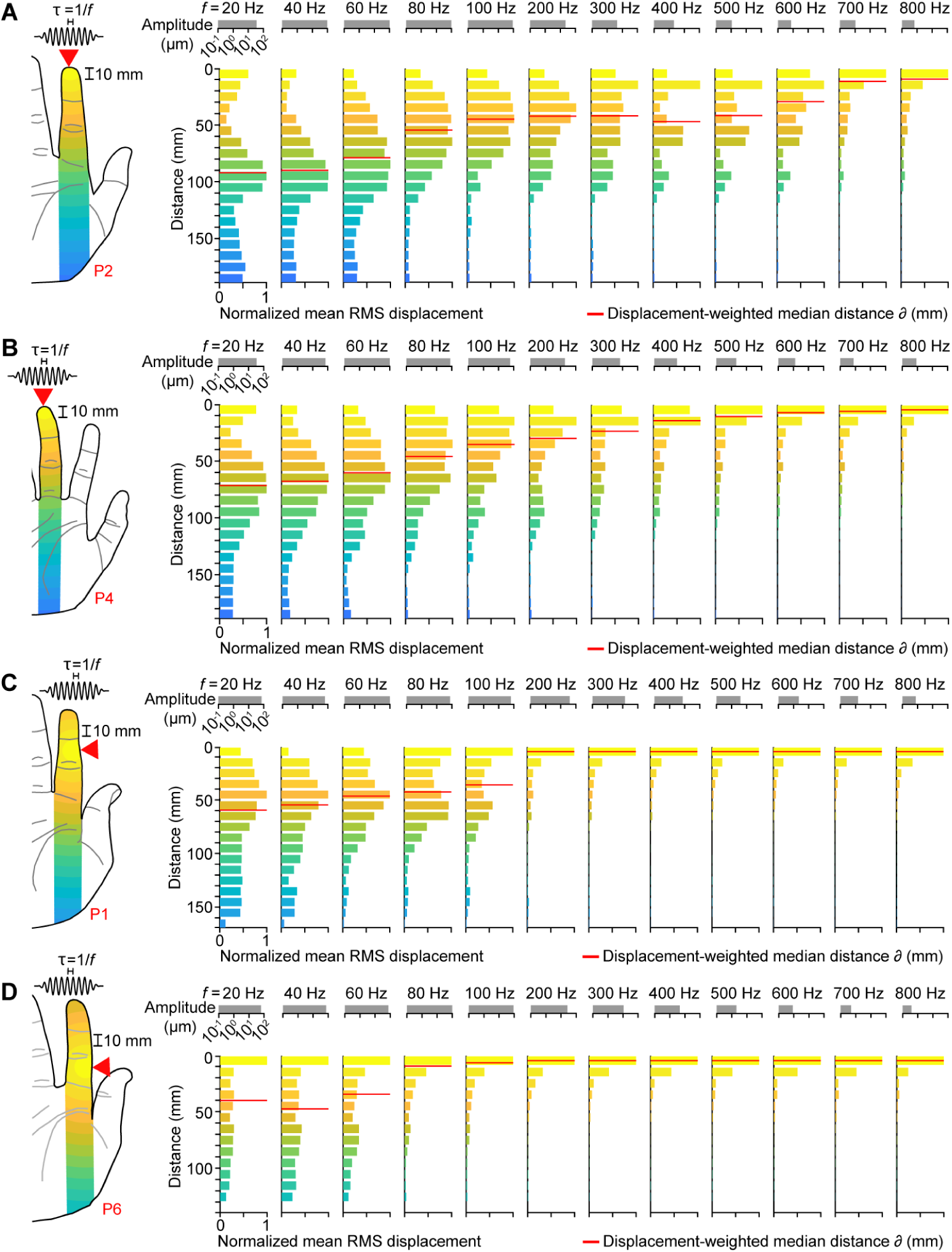
RMS skin displacement distributions across frequency for other contact locations and participants. (A) Normalized root mean square (RMS) skin displacements averaged within 10 mm-wide bands at increasing distances from the contact location (digit II DP) elicited by sinusoidal stimuli of various frequencies (20 to 800 Hz) for P2. Amplitude scale bars (top, gray) show maximum peak-to-peak displacement across the hand at each frequency. Horizontal red lines: median transmission distance; red text: participant number; red arrow: contact location. (B) As in (A) for P4 and stimuli applied at the digit III DP. (C) As in (A) for P1 and stimuli applied at the digit II IP. (D) As in (A) for P6 and stimuli applied at the digit II PP.

**Figure S4.**
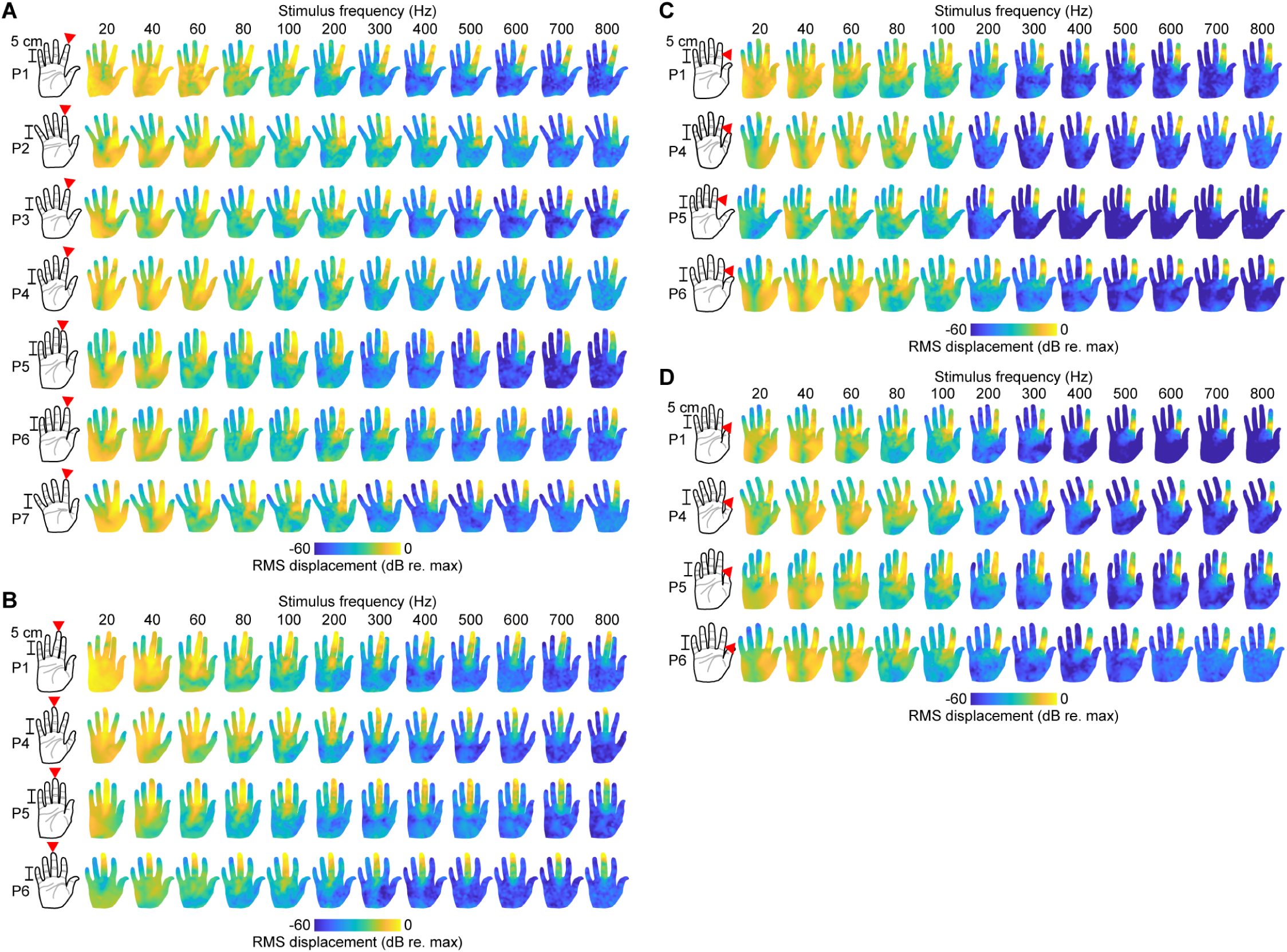
Whole-hand RMS skin displacements across frequency for all contact locations and participants. (A) Normalized whole-hand RMS skin displacements (log scale) elicited by sinusoidal stimuli of various frequencies (20 to 800 Hz) applied at the digit II DP. Shown for all participants. Red text: participant number; red arrow: contact location. (B) As in (A) for stimuli applied at the digit III DP. (C) As in (A) for stimuli applied at the digit II IP. (D) As in (A) for stimuli applied at the digit II PP.

**Figure S5.**
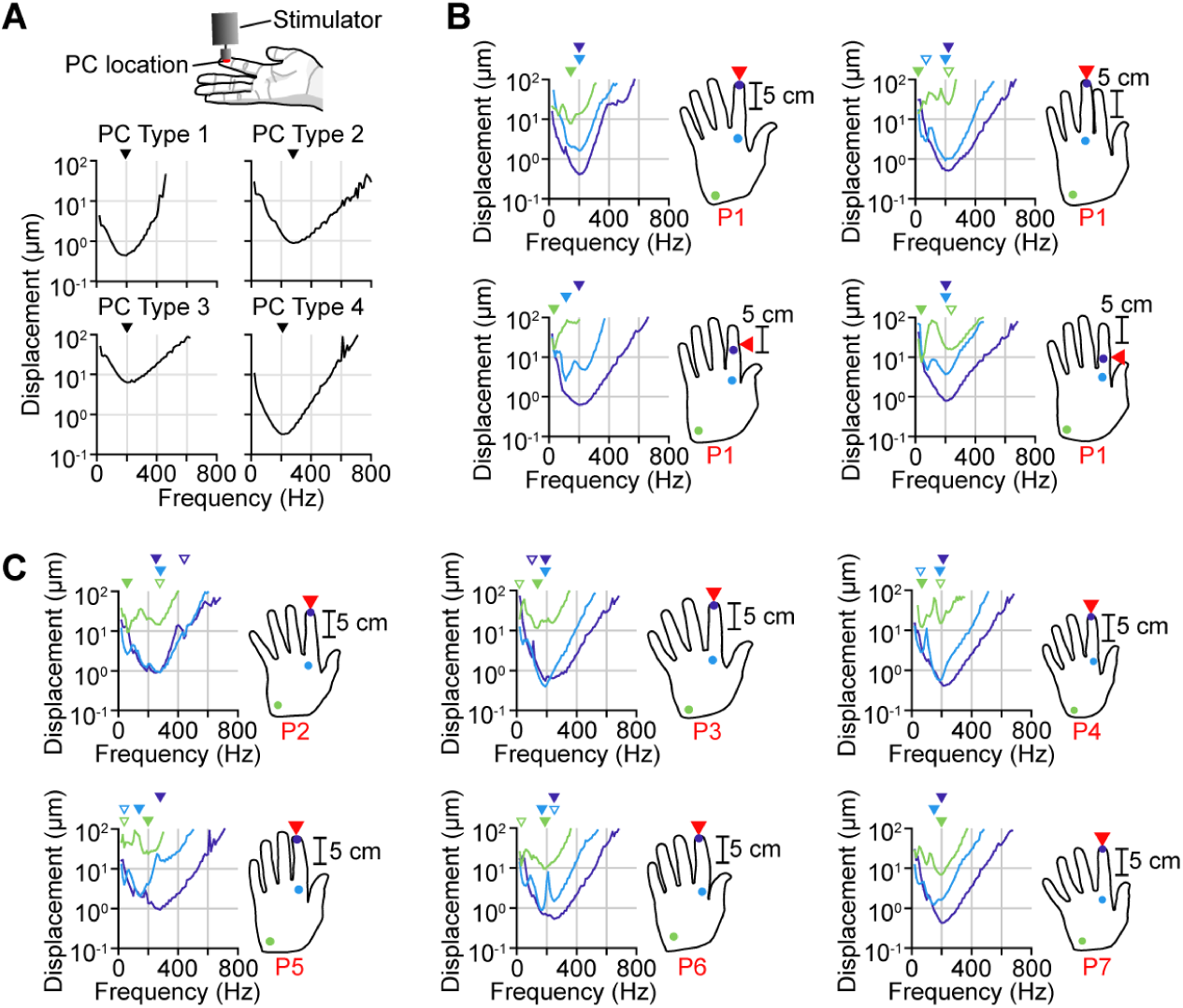
PC entrainment threshold curves for other contact locations and participants. (A) Entrainment threshold curves for PCs located directly underneath the contact location. Shown for all four types of PC neuron models. Triangles above curves: global minimum. (B) Entrainment threshold curves for PCs at three locations on the hand for stimuli applied at four different contact locations. Shown for PC model type 4. Red arrow: contact location; red text: participant number; color: PC locations; filled triangles above curves: global minimums; unfilled triangles above curves: other local minima (prominence > 6 dB). (C) As in (B), for other participants.

**Figure S6.**
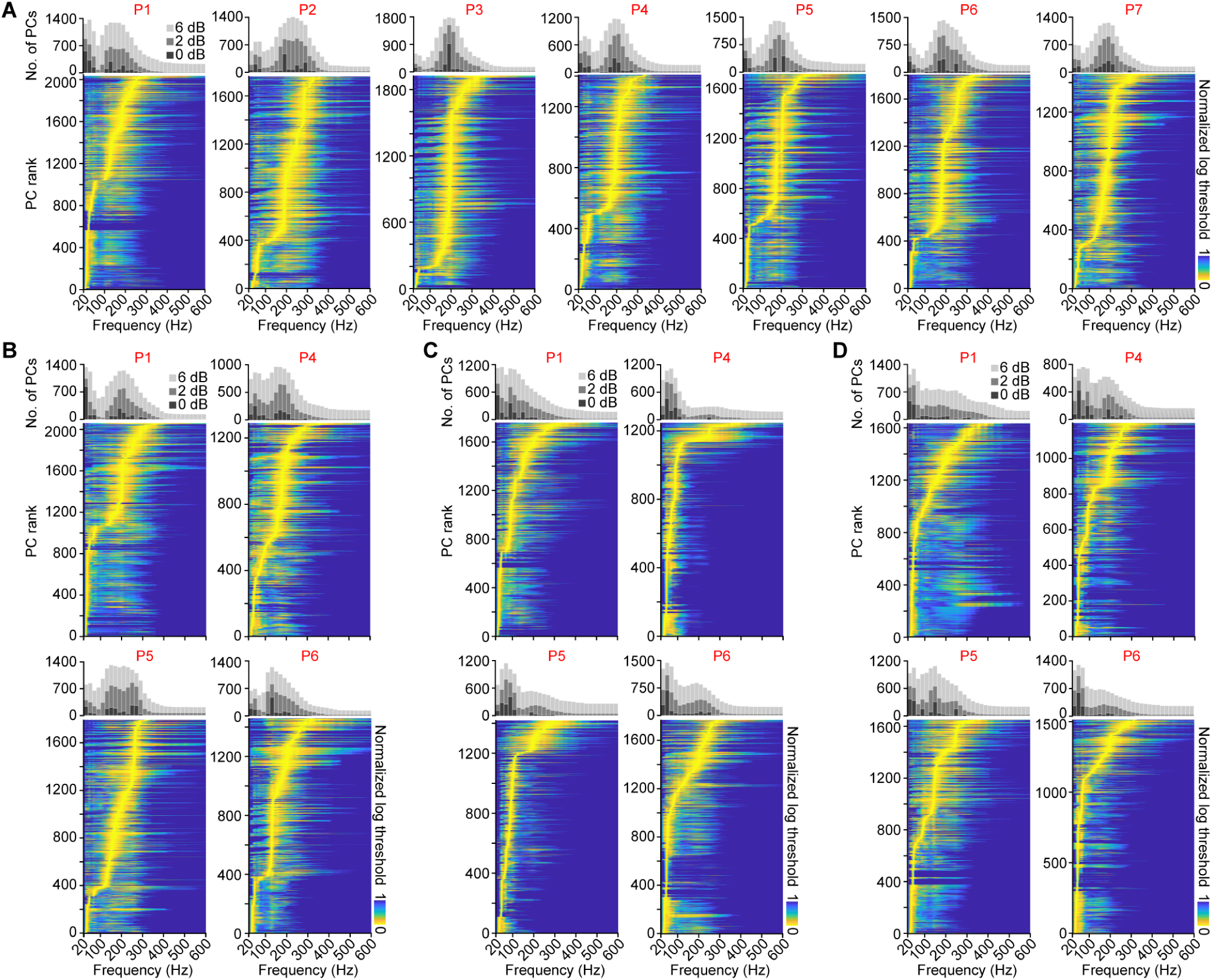
Rank-ordered PC entrainment threshold curves for all contact locations and participants. (A) Entrainment threshold curves for all PCs in the whole-hand population rank-ordered by preferred frequency for stimuli applied at the digit II DP. Each curve is shown on a log scale and normalized from 0 to 1. Histograms show the number of PCs at each frequency with entrainment threshold curve values within 0 dB (light gray), +2 dB (medium gray), and +6 dB (dark gray) of the minimum. Red text: participant number. (B) As in (A), for stimuli applied at the digit III DP. (C) As in (A), for stimuli applied at the digit II IP. (D) As in (A), for stimuli applied at the digit II PP.

**Figure S7.**
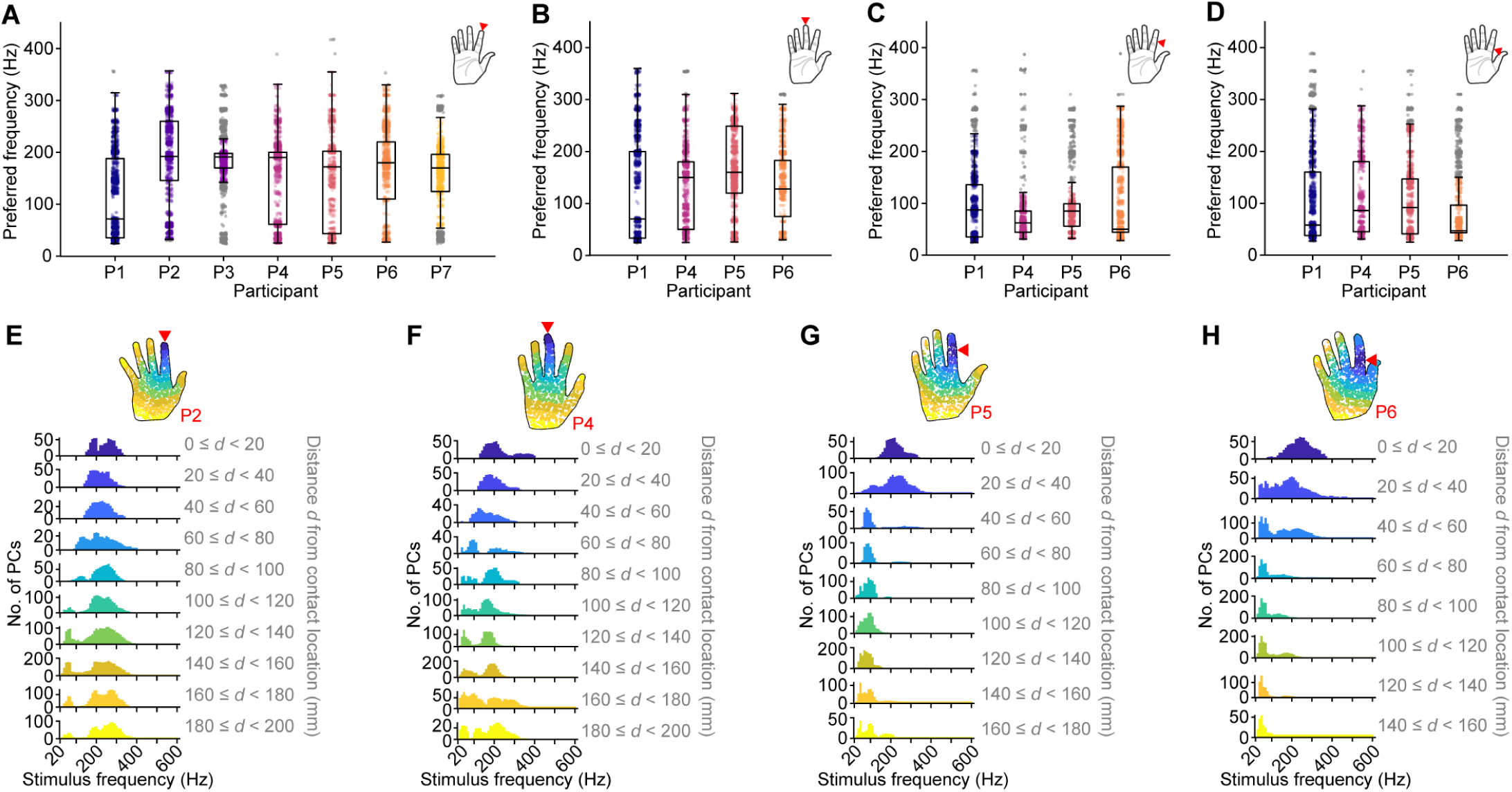
Frequency sensitivity in whole-hand PC populations. (A) Preferred entrainment frequencies across whole-hand PC populations for each participant for stimuli applied at the digit II DP. Red arrow in hand plot (top right): contact location; lower box limits: 25th percentile; upper box limits: 75th percentile; lines within boxes: 50th percentile; whiskers: range of data within one IQR from the lower or upper box limits; colored dots: preferred frequencies within the whiskers; gray dots: outliers. (B) As in (A), for stimuli applied at the digit III DP. (C) As in (A), for stimuli applied at the digit II IP. (D) As in (A), for stimuli applied at the digit II PP. (E) Histograms summarizing the number of PCs at each frequency with entrainment threshold curve values within +2 dB of the global minimum for stimuli applied at the digit II DP. PC subpopulations corresponding to each histogram are constructed by selecting PCs located within 20 mm-wide bands at increasing distances from the contact location. Histogram color corresponds to the colored PC subpopulations shown in the hand plot (top). Gray text: distance d from contact location; red arrow: contact location; red text: participant number. (F) As in (E), for stimuli applied at the digit III DP. (G) As in (E), for stimuli applied at the digit II IP. (H) As in (E), for stimuli applied at the digit II PP.

**Figure S8.**
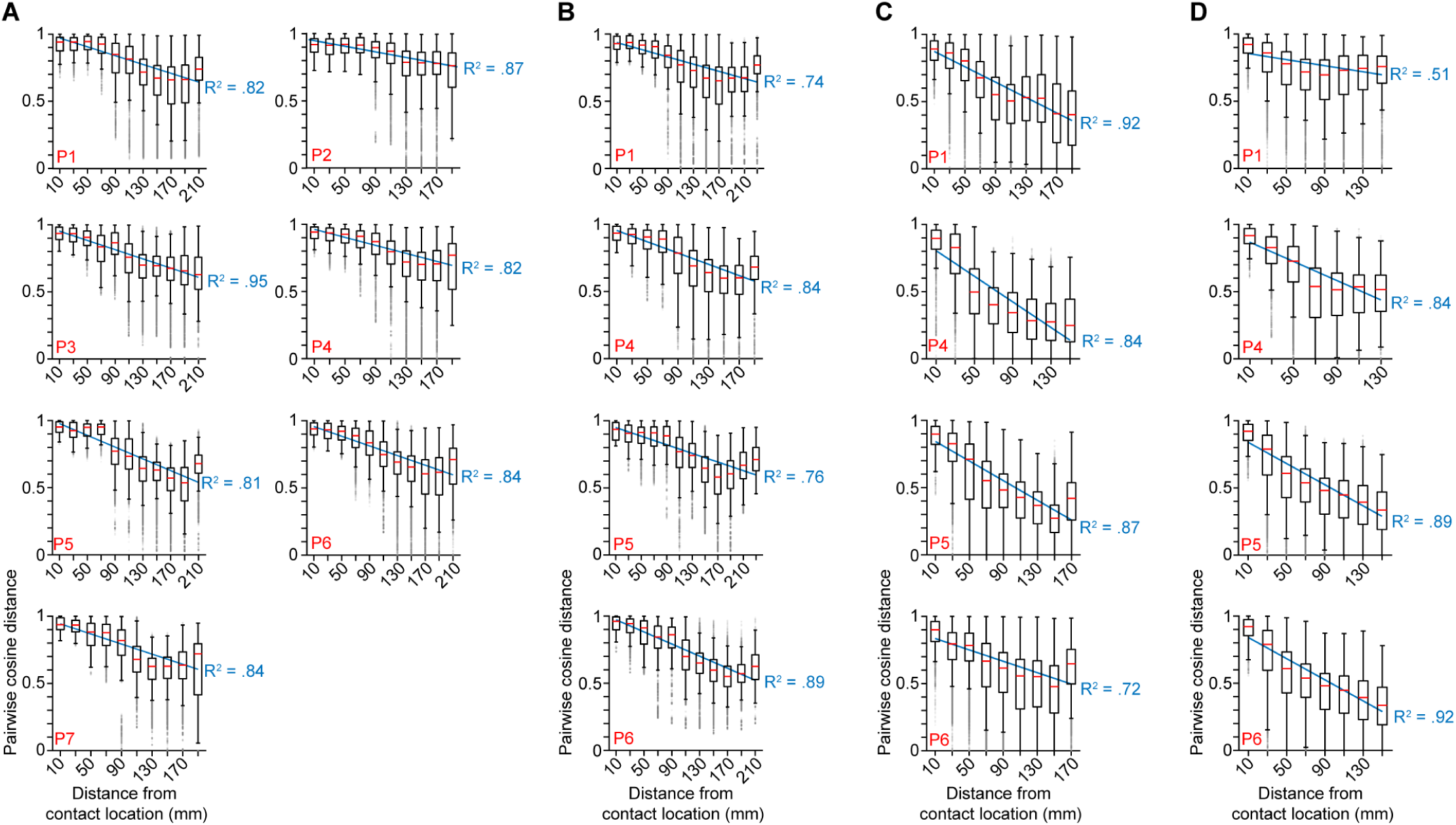
Entrainment threshold curve cosine distances between PC subpopulations at various distances from the contact location. (A) Cosine distances between pairs of entrainment threshold curves of PCs located within 10 mm of the contact location (digit II DP) and those of PCs located within 20 mm-wide bands at increasing distances from the contact location. X-axis labels denote the center distance of each band. Red midlines: median; lower box limits: 25th percentile; upper box limits: 75th percentile; whiskers: range of data within one IQR from the lower or upper box limits; gray dots: outliers; gray dotted lines: linear fit of medians; gray text: R^2^ value for linear fit; red text: participant number. (B) As in (A), for stimuli applied at the digit II IP. (C) As in (A), for stimuli applied at the digit II IP. (D) As in (A), for stimuli applied at the digit II PP.

**Figure S9.**
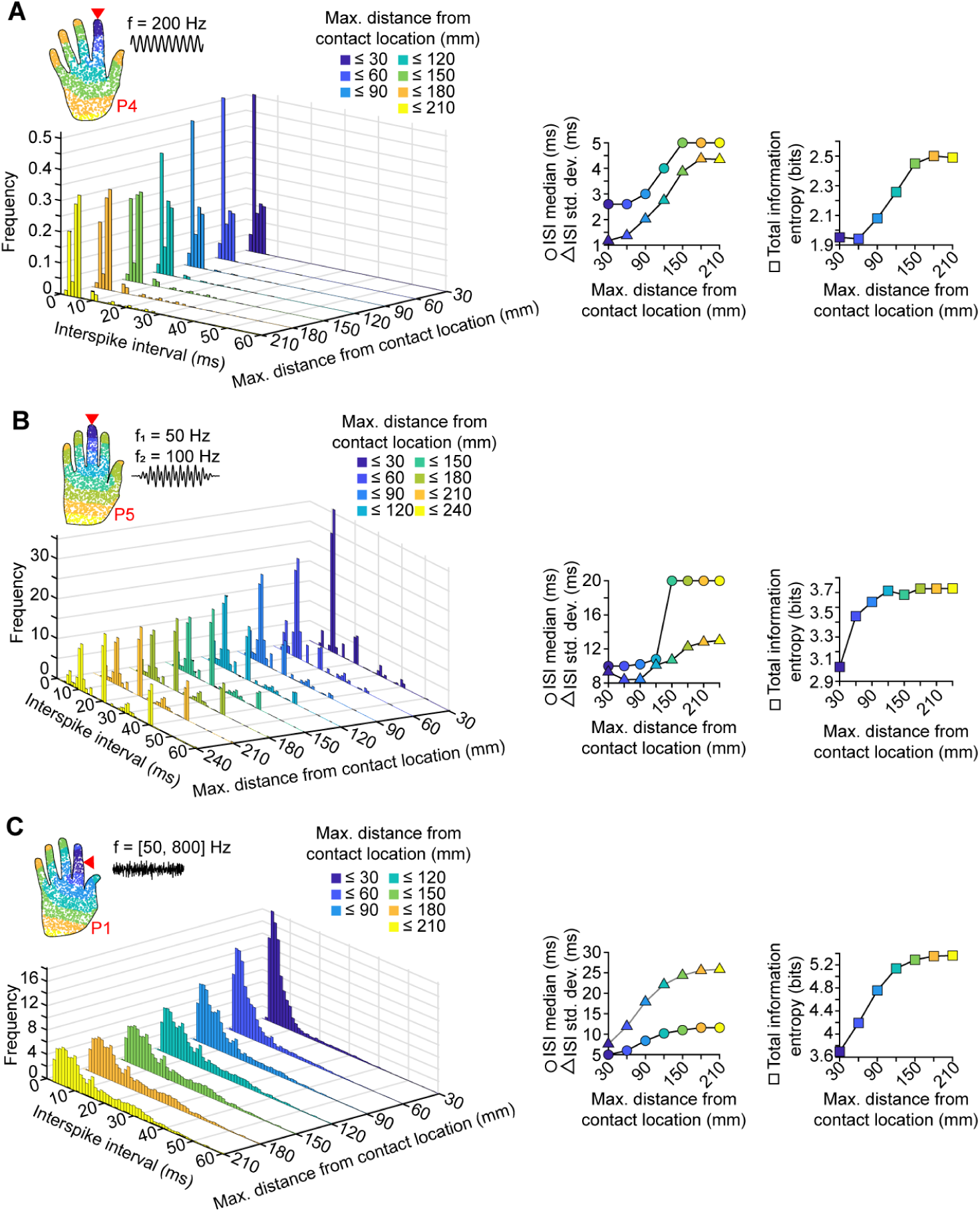
ISI distributions constructed from PCs located within increasing distances from the contact location. (A) Left panel: Histograms comprising interspike intervals (ISIs) from PCs located within increasing distances from the contact location (hand inset) in response to a sinusoidal stimulus (200 Hz, 15 µm max. peak-to-peak displacement across hand). Right panel: median (circles), standard deviation (triangles), and total information entropy (squares) of the ISI histograms shown above. Color: maximum distance from the contact location; red arrow: contact location; red text: participant number. (B) As in (A), for a diharmonic stimulus (f_1_ = 50 Hz, f_2_ = 100 Hz, 10 µm max. peak-to-peak displacement across hand for both f_1_ and f_2_). (C) As in (A), for a bandpass-filtered noise stimulus (50 to 800 Hz band, 5 µm max. RMS displacement across hand).

**Figure S10.**
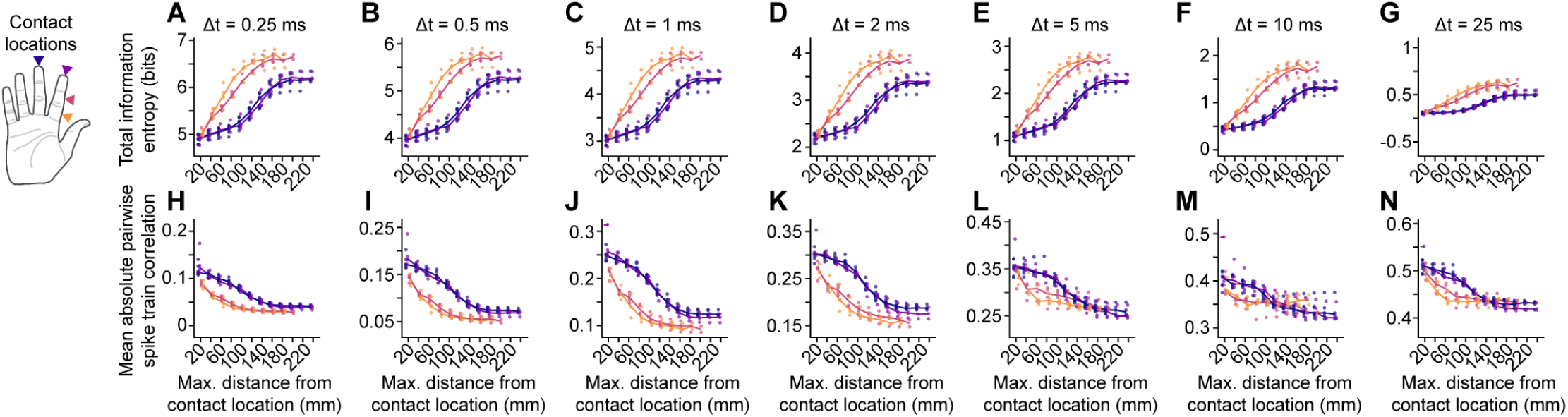
Information entropy of ISI histograms and mean spike train correlations for different bin sizes. (A) Total information entropy (bits) of histograms comprising ISIs from PCs located within increasing distances from the contact location in response to a diverse stimulus set (see Methods) for an ISI histogram bin width of Δt = 0.25 ms. Colors: contact location corresponding to arrows in hand plot (top left); dots: data points for all participants; lines: median. (B) As in (A), for an ISI histogram bin width of Δt = 0.5 ms. (C) As in (A), for an ISI histogram bin width of Δt = 1 ms. (D) As in (A), for an ISI histogram bin width of Δt = 2 ms. (E) As in (A), for an ISI histogram bin width of Δt = 5 ms. (F) As in (A), for an ISI histogram bin width of Δt = 10 ms. (G) As in (A), for an ISI histogram bin width of Δt = 25 ms. (H) Mean absolute spike train correlation between all pairs of PCs located within increasing distances from the contact location for a spike train histogram bin width of Δt = 0.25 ms. Colors: contact location corresponding to arrows in hand plot (top left); dots: data points for all participants; lines: median. (I) As in (H), for a spike train histogram bin width of Δt = 0.5 ms. (J) As in (H), for a spike train histogram bin width of Δt = 1 ms. (K) As in (H), for a spike train histogram bin width of Δt = 2 ms. (L) As in (H), for a spike train histogram bin width of Δt = 5 ms. (M) As in (H), for a spike train histogram bin width of Δt = 10 ms. (N) As in (H), for a spike train histogram bin width of Δt = 25 ms.

## Supplemental Tables

**Table S1.** Sinusoidal stimulus set parameters for PC population spiking activity analysis.

| Frequency (Hz) | D<sub>pp</sub> amplitudes (μm) |
| --- | --- |
| 50 | 2.3, 5.9, 15.2, 38.9, 100 |
| 75 | 1.6, 4.5, 12.7, 35.6, 100 |
| 100 | 1.1, 3.4, 10.5, 32.4, 100 |
| 150 | 0.5, 1.9, 7.1, 26.6, 100 |
| 200 | 0.3, 1.4, 5.8, 24.2, 100 |
| 250 | 0.4, 1.4, 5.9, 24.3, 100 |
| 300 | 0.6, 2.2, 7.8, 28.2, 100 |
| 400 | 2.1, 5.2, 13, 32.2, 80 |
| 500 | 2.8, 5.8, 11.8, 24.3, 50 |
| 600 | 7.3, 10.8, 16, 23.7, 35 |
| 700 | 21, 21.9, 23, 24, 25 |
| 800 | 10, 11.9, 14.1, 16.8, 20 |

**Table S2.**
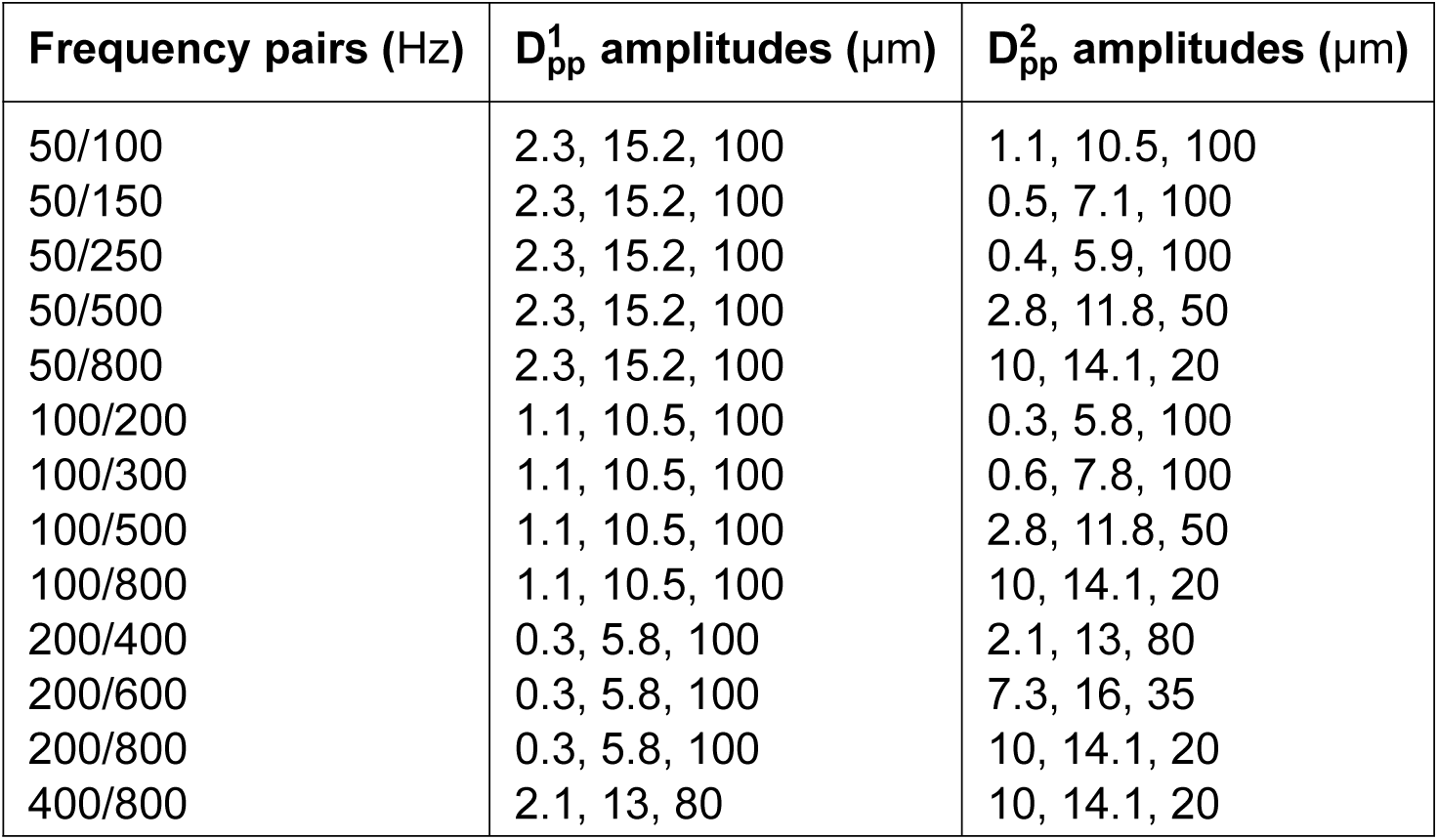
Diharmonic stimulus set parameters for PC population spiking activity analysis.

**Table S3.** Bandpass noise stimulus set parameters for PC population spiking activity analysis.

| **Frequency band (Hz)** | **D<sub>RMS</sub> amplitudes (μm)** |
| --- | --- |
| 50 to 100 | 0.5, 1, 5, 10, 25 |
| 50 to 250 | 0.5, 1, 5, 10, 25 |
| 50 to 500 | 0.5, 1, 5, 10, 25 |
| 50 to 800 | 0.5, 1, 5, 10, 25 |
| 100 to 250 | 0.5, 1, 5, 10, 25 |
| 100 to 500 | 0.5, 1, 5, 10, 25 |
| 100 to 800 | 0.5, 1, 5, 10, 25 |
| 250 to 500 | 0.5, 1, 5, 10, 25 |
| 250 to 800 | 0.5, 1, 5, 10, 25 |
| 400 to 800 | 0.5, 1, 5, 10, 25 |

